# A single dynamical property can account for the capacity to learn, from artificial networks to the mammalian brain

**DOI:** 10.64898/2026.07.09.737603

**Authors:** Ravi Chopra, Juntao Zhong, Ezra Miller, Gemechu Bekele Tolossa, Leandro J. Fosque, Jordan Janzen-Meza, Nicholas W. DeKorver, Rejean Guerriero, Neil J. Ritter, Mary E. Lambo, Kiran Bhaskaran-Nair, Stephen Van Hooser, Woodrow Shew, Keith B. Hengen

## Abstract

Every brain must adapt to an unpredictable world, yet individuals differ in how readily they learn. Theoretical work suggests that learning is fastest when a system, whether biological or synthetic, is initialized in a state close to instability - i.e., near *criticality* - because critical dynamics are imbued with a diverse repertoire of patterns and multi-scale correlations. Here, we empirically estimate distance to criticality in the brain and show that it predicts the rate of adaptability underlying learning, neuronal tuning, and general intelligence. In mouse motor cortex, proximity to criticality forecasts the learning rate of two future, complex tasks: prey-capture hunting and ladder crossing. In contrast, distance to criticality predicted neither an animal’s naive ability nor its asymptotic skill — isolating the rate of learning itself. In visual cortex of young ferrets, proximity to criticality predicts how strongly subsequent experience reshapes neural tuning. In human frontal cortex, it correlates with general cognitive ability. A minimal recurrent network model reproduced these results and offers a mechanism: proximity to criticality defines the timescale over which a system can learn from its past experiences, directly setting the rate of learning.

## Introduction

What is the basis of an individual’s durable intelligence? What is the biological foundation of the capacity to learn? Any complex system capable of adapting to an unpredictable environment should benefit from exploring as large a space of possible solutions as quickly as possible. This principle is well established in artificial intelligence and machine learning, where solutions are found quickly only when diverse patterns of activation spread across nodes and layers without saturating the system^1–5^. Does learning in biological systems obey the same principle? If so, a brain that generates a larger repertoire of activity patterns should be the more adaptable and thus learn faster.

How should brain dynamics be configured to achieve such an adaptable state? Converging insights from machine learning, neuroscience, and physics point to a compelling answer. In machine learning, learning proceeds fastest when a network is initialized near the boundary between two dynamical regimes — stability and chaos — a state variously termed the “edge of chaos”^6–9^ or “criticality”^1,4,10–13^. This holds whether learning is driven by conventional gradient-descent backpropagation^1,3–5,14^ or by more biologically realistic rules^2^. Moreover, keeping a network near criticality throughout learning, rather than only at initialization, further improves the quality of what it learns^3,14^. This marginally stable, critical state is what endows the system with the large repertoire of activity patterns thought to accelerate learning^15–18^. The reason is rooted in the fundamental physics of criticality: by renormalization group theory^19,20^, scale invariance in interacting systems arises *exclusively* at critical fixed points — other arrangements of a system generate correlations that disappear across scales^21^. It is therefore only at criticality that a system can generate patterns of every scale, from small to large and fleeting to persistent. This informs our central hypothesis: living organisms learn faster the closer they are to criticality.

Congruent with a role in the brain’s general ability to produce adaptive behavior, near critical dynamics are evident across species and are consistent with an endpoint of homeostatic plasticity^10,22–26^. As biology cannot tune to a point, homeostatically-controlled physiologic parameters, such as body temperature and blood pH^27^, are actively maintained within a viable range. This must also be true of similar set-points in brain dynamics. For this reason, a brain cannot “be critical”; instead, neuronal activity should be described according to its *distance* from a critical boundary^28^. As a population is tuned further and further from criticality, the scale-invariant range shrinks^29,30^, and computational capacity diminishes^16,31–34^. Additionally, when considering homeostatically controlled physiological features, population-level variance is often larger than individual-level variance. Plainly, there is truth to the idea that some people run hotter than others^35^. It stands to reason that, as a homeostatic set-point, proximity to criticality should be no different. Specifically, should healthy brain circuits be actively maintained near criticality^22,23^, some circuits and/or individuals may be expected to be consistently closer (or further) than others. If true, and if proximity to criticality establishes the capacity for adaptive computation, then proximity to criticality should both determine the facility with which a brain can learn, and account for persistent inter-individual differences in the rate of learning and adaptation.

## Results

### Isocortical proximity to criticality predicts future learning

To directly test the hypothesis that baseline feature(s) of an animal’s neural dynamics account for inter-individual differences in learning (Fig. 1A), we continuously tracked the activity of ensembles of putative excitatory single neurons (regular spiking units; RSUs^36–38^) in right primary motor cortex (M1) of adult wild-type C57BL/6 mice (*N* = 18) for a total of 13 to 14 days (312–336 h) per animal (6,326 RSUs total across all blocks, mean = 34 ± 13.9 per block, minimum = 15 per block, maximum = 71 per block) (Fig. 1B). This 13–14 d continuous recording comprised a 24–48 h baseline of spontaneous behavior in the home cage, followed by 6 d in each of two motor learning tasks, prey capture and ladder crossing (Fig. S1). Animals were randomly assigned to ladder first or prey capture first. All animals included in these experiments were wild type (WT) and thus expected to exhibit modest variability in cognitive performance relative to diseased animals or a genetically heterogeneous population. We attempted to forecast future learning in each task from a suite of neural and behavioral measures collected before animals had any experience with either paradigm (Fig. 1C,D). During the baseline period, three arousal states were labeled based on standard polysomnography (local field potential spectral power, video)^23,39^: wake, NREM sleep, and REM sleep. Here, unless otherwise mentioned, all neural features are examined exclusively during waking epochs.

**Figure 1:**
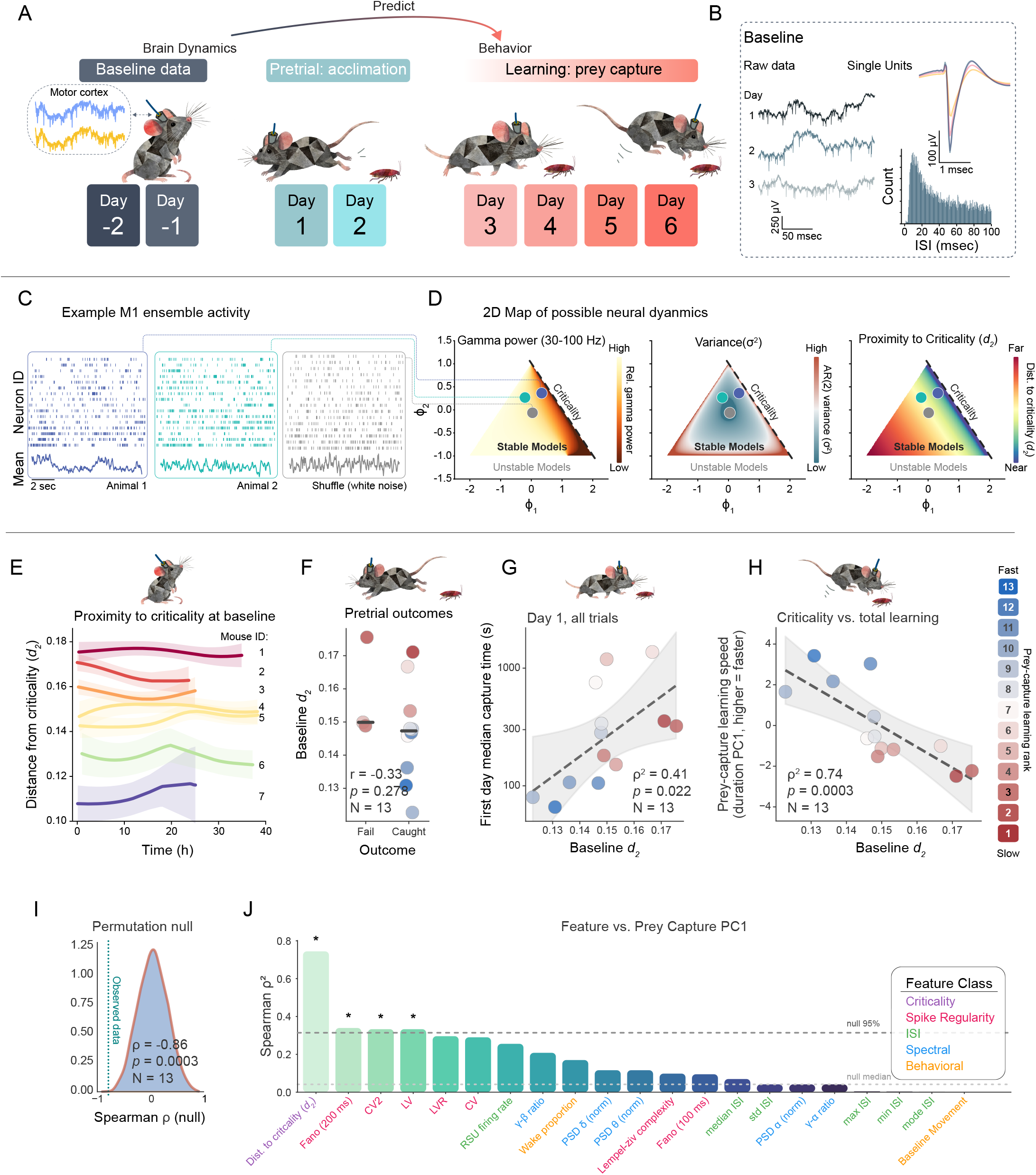
Proximity to criticality predicts prey-capture learning rate. **(A,B)** Experimental design: ensembles of single units in primary motor cortex were recorded for two baseline days. Baseline, single unit neural activity was used to predict various aspects of a subsequent six day prey-capture learning protocol. Pretrial acclimation allowed mice to acclimate to cockroaches. Next, mice completed 24 food-deprived hunting trials spread across four days. **(C)** Example ensemble rasters and population activity range from richly structured near criticality (low *d*_2_) to progressively flatter and more noise-like as *d*_2_ grows, with a spike- time-shuffled control approximating white noise. **(D)** Autoregressive (AR) models reproduce physiologically- relevant neural dynamics. A second-order AR model is defined by two variables, *ϕ*_1_ and *ϕ*_2_, allowing a 2D visualization of all possible models. The central triangle comprises the set of models with stable dynamics. These can be quantified in terms of, e.g., gamma band power **(left)**, variance **(center)**, and the shortest distance to a critical model **(right)**. Note that, in this 2D space, the upper right boundary of stable models (dashed line) is the set of models that generate scale-invariant, critical activity. **(E)** At baseline, proximity to criticality (measured as *d*_2_ during waking epochs) is stable within individual animals over time. **(F)** Baseline *d*_2_ does not predict whether or not mice consume the pretrial cockroach (*r* = 0.33*, p* = 0.278). **(G)** In contrast, on the first day of trials (i.e., the first opportunity to refine behavior as a function of experience), baseline *d*_2_ predicts median capture time - where mice closer to criticality at baseline are faster hunters (*ρ*^2^ = 0.41*, p_perm_* = 0.022, dashed line, fit; shaded band, 95% CI). **(H)** Across all 24 trials over four days, *d*_2_ at baseline predicts a four-measure composite of prey-capture learning (*ρ*^2^ = 0.74*, p_perm_* = 0.0003, dashed line, fit; shaded band, 95% CI). F-G report the *N* = 13 mice that hunted. Individuals are colored by their learning ranking (H, right). **(I)** A permutation null confirms that the learning effect lies outside chance. Observed value, dashed line; *p_perm_* = 0.0003). **(J)** The predictive value of twenty one features measured during the baseline. Features include neural, spectral, and behavioral metrics. In addition to *d*_2_, three spike-regularity measures (Fano-200 ms, CV2, LV) clear the 95th-percentile permutation null (dashed line; asterisks).

Any sample of neural data can be described in terms of a wide range of metrics, including firing rate, variance, spectral power, and more. Each of these measures is likely related to the tendency to generate the diverse repertoire of activity patterns needed for fast learning. However, a direct test of our hypothesis requires a direct measurement of proximity to criticality. For this, we use *d*_2_, a recently developed method for empirically estimating proximity to criticality based on univariate time series^40^. By applying renormalization group theory to autoregressive models, *d*_2_ quantifies the extent to which a given sample reflects scale-invariant fluctuations (Fig. 1D, S2). Concretely, purely sinusoidal activity is defined by a single time scale, while critical dynamics are defined by correlations at all scales. In practice, *d*_2_ is calculated by first fitting a model to observed data and then measuring the distance from the best fit model to the nearest model whose activity is scale invariant (i.e., distance to the dashed border in Fig. 1D). Briefly, we converted the spike times of single unit ensembles into a population spike count timeseries. We fit a 10^th^ order autoregressive (AR) model to the time series, yielding 10 AR coefficients, thus specifying a point in a 10-dimensional parameter space of all possible 10^th^ order AR models (example for AR(2) model in Fig. 1D). In this space, *d*_2_ is the minimum Euclidean distance between the best-fit model point and the hyperplane formed by the set of critical models, i.e., 10^th^ order models that generate autocorrelations across all scales. In our animals, the mean motor cortex *d*_2_ was 0.1494 ± 0.000489 (*N* = 18). For comparison, we shuffled spike times to maintain population activity rate while destroying temporal correlations, which provides a quantitative example of *d*_2_ for dynamics ‘very far’ from criticality (Fig. 1C). This produced mean *d*_2_ *_shuffle_* of 0.3166±0.00006, close to the analytical solution for white noise with order 10 (*d* for 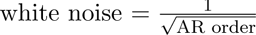) of *d*_2_ = 0.32. Practically, white noise defines the upper limit of *d*_2_ short of invoking an unstable regime (for example, seizure). Note that a wide range of physiologically-plausible dynamics are well described by AR models — the same AR landscape can thus be evaluated in terms of any descriptor of neural activity; Fig. 1D shows the same AR(2) landscape as a function of gamma power, variance, and *d*_2_. This offers a direct comparison of alignment across metrics (Fig. 1D). Any movement in this landscape invariably entails changes in multiple features.

### Isocortical d_2_ reliably differentiates animals

If distance from criticality is homeostatically regulated, it is, by definition, actively maintained within some effective range^22^. While animals could, in theory, drift without constraint within this range, many homeostatically controlled parameters show comparatively tight control at the level of the individual organism^37,41–43^. As a result, an individual may be typified as “running hot” relative to another. Whether distance to criticality can reliably identify an animal is unknown, yet this is a logical predicate for our hypothesis that it comprises a durable feature of cognition. To assess this, we examined the source of variance in *d*_2_ during the baseline (Fig. 1E). Considered across *N* = 18 animals, baseline *d*_2_ spanned a 1.6-fold range of setpoints (per-animal means 0.111 − 0.176; between-animal *SD* = 0.015) (Fig. 1F). Individual 5-min estimates of *d*_2_ were themselves noisy, such that animal identity accounted for only ∼ 30% of the variance in single windows (one-way random-effects *ICC*(1, 1) = 0.30, 95% CI 0.15 − 0.42), the remainder reflecting fast within-animal fluctuation (Fig. S3). This within-animal variance, however, largely averaged out (*ICC*(1*, k*) = 0.94). Each animal’s setpoint (baseline mean) was a stable and reproducible feature. Between-animal differences in setpoint exceeded chance even after accounting for the autocorrelation of successive windows (one-way ANOVA on independent 3 h blocks, *F* (17, 79) = 12.6, *p_perm_*= 1 × 10*^−^*^4^ by label permutation). Furthermore, an animal’s setpoint estimated from the first half of its baseline predicted the setpoint measured over the second, non-overlapping half (test– retest *r* = 0.83, 95% CI 0.58 − 0.94; Spearman–Brown full-length reliability = 0.91). Consistent with a durable individual trait, baseline *d*_2_ alone was sufficient to distinguish animals (Fig. S3). Pairwise discriminability rose monotonically with the difference in setpoint (Spearman *ρ* = 0.94 between AUC and setpoint separation, *p <* 10*^−^*^69^), reaching a median AUC of 0.68 from a single 5-min window and 0.89 with ∼ 2.7 h of data, and a cross-validated classifier identified the animal from *d*_2_ at twice the chance rate across all 18 individuals (*p_perm_* = 5 × 10*^−^*^4^). Thus, although distance to criticality fluctuates from moment to moment, its setpoint is a reliable individual trait. Some animals sit consistently closer to criticality than others.

In the five animals implanted in both M1 and V1, simultaneously recorded *d*_2_ shared only a modest fraction of its moment-to-moment variance across the two regions (Spearman *ρ* = 0.33 for *d*_2_ as predictor of per-animal median) and held distinct regional set-points, indicating that proximity to criticality is regulated locally within a circuit rather than imposed by a single brain-wide metabolic or global physiological variable (Fig. S4).

### Isocortical proximity to criticality forecasts prey-capture learning rates

Mathematically, a system’s proximity to criticality constrains the transmission of information through time and across space^31^. Thus, given a fixed number of neurons, distance from criticality should directly limit the extent and rate of learning. On this basis, we reasoned that animals tuned closer to criticality should learn more quickly than those tuned further. To test this, we asked whether animal-level set-points in proximity to criticality during the baseline period could forecast learning rate in either of two six-day tasks: prey capture and ladder crossing. We first examined prey capture, in which mice that were bred and raised in an academic vivarium were introduced tored runner cockroaches (*Shelfordella lateralis*). All mice were acclimated to red runners during an overnight pre-trial exposure (days 1–2) with *ad lib* food access, followed by food-deprived hunting trials (days 3–6). Five of 18 mice never hunted, and were thus excluded from these analyses, and whether a mouse hunted or not was independent of *d*_2_ (S5A).

Importantly, learning is the means by which an animal transitions from its starting point to its end point via experience. In the context of our prey-capture task, whether an animal is initially frightened or eventually athletic should be largely independent of the intervening period of plastic refinement. Consistent with this, in wild type animals, distance to criticality at baseline carried no information about how animals responded to the pretrial red runner. Note that, during the pre-trial, mice acclimated to a novel animate object, overcoming neophobia and, over hours, most slowly realized the cockroach could be treated as a food source. Of the hunters (13/18), 77% (10/13) caught and consumed the pretrial red runner, which was unrelated to baseline *d*_2_ (point-biserial *r* = −0.33, *p* = 0.28; Fig. 1F). Pre-trial capture time ranged from 0.12 to 11.8 h and was also not predicted by baseline *d*_2_ (*ρ*^2^ = 0.006, *p_perm_*= 0.83, Fig. S5B). Thirty six hours after the pre-trial night, food-deprived hunting trials began. Mice completed a total of 24 hunting trials (six red runners / day for four days). The first hunting trial is unique in that it arises with a motivation to pursue food but no prior hunting experience, and thus reflects an animal’s starting point for learning. In line with our hypotheses, murine proximity to criticality did not predict performance on the first trial (*ρ*^2^ = 0.143, *p_perm_*= 0.23; Fig. S5C), suggesting that the dynamical regime of M1 indicates nothing about innate hunting prowess or motivation. Following the first trial, all animals exhibited robust improvements in hunting performance even on Day 1: every hunter (13/13) was faster on a subsequent Day-1 trial than on its naive first trial (Fig. S5D). Median capture time dropped from 761 (Trial 1) to 95 *s* (fastest capture time on Day 1) across mice, an 88% reduction (Wilcoxon signed-rank *p* = 0.000244).

When considering the first opportunity for learning (Day 1 trials), mice closer to criticality at baseline were more effective in learning to hunt quickly than those further from criticality. In contrast to the very first trial, baseline *d*_2_ predicted Day 1 median capture time (*ρ*^2^ = 0.41, *p_perm_* = 0.02, Fig. 1G). All animals continued to shorten capture time across the 24 trials (13/13 with a negative trial-vs-time correlation, median Spearman *ρ* = −0.70, sign-test *p* = 1.2 × 10*^−^*^4^), with gains decelerating toward an asymptote within the first ∼10–15 trials (exponential *τ* ≈ 6 trials; day-over-day reduction falling from 223 s to 14 s, paired Wilcoxon *p* = 5 × 10*^−^*^4^) (Fig. S5D). To avoid over-reliance on any one metric, we quantified learning along four axes related to two independent methods (breakpoint analysis and exponential fitting of capture times, Fig. S5E-G). Individually, the four metrics were highly concordant (Kendall’s *W* = 0.88, Fig. S5F), and we constructed a composite learning-rate metric as the first principal component across the four measures, which captured 89% of their shared variance. Considered over the four day, 24 trial protocol, baseline proximity to criticality was strongly correlated with this composite (Spearman *ρ*^2^ = 0.74, *p*_perm_ = 0.0003, *N* = 13; Fig. 1H): mice closer to criticality at baseline — i.e. with lower *d*_2_ — acquired the skill more rapidly, with baseline *d*_2_ alone explaining ∼ 74% of the variance in learning rate. This relationship did not depend on any single animal (leave-one-out Spearman *ρ* ∈ [−0.888, −0.832]), and each constituent metric tracked baseline *d*_2_ on its own (Fig. S5G). We quantify the *d*_2_–learning relationship with Spearman’s *ρ* because hunting performance is a bounded, saturating outcome. Once learning is complete (i.e., animals have reached their behavioral asymptote), *d*_2_ was meaningless as a predictor: the median time-to-subdue over each animal’s final ten trials (22–220 s across mice) did not covary with baseline *d*_2_ (Spearman *ρ* = +0.36, *p_perm_* = 0.23), nor did the fitted asymptote of the capture-time learning curve (*ρ* = −0.29, *p_perm_* = 0.34), nor the mean late-phase capture latency (*ρ* = +0.24, *p_perm_* = 0.44) (Fig. S6A-B). Baseline *d*_2_ likewise carried no signature of athletic capacity: mean, peak, and total speed during hunts did not significantly covary with distance to criticality (all *p* ≥ 0.08; data not shown). These findings, paired with our findings during the learning phase, reinforce the conclusion that proximity to criticality specifically predicts rate of experience-dependent improvement in hunting performance, without any relationship to the point where learning started or its ultimate result: in the vocabulary of machine learning, a learning rate.

While the strength of this relationship, particularly in a freely behaving neurobiological context, is perhaps surprising, we reasoned that it may not be unique^44,45^. To assess this, we analyzed the predictive power of 20 additional neural, spectral, and behavioral features from the baseline period (Fig. 1J). Only 3/20 features rose above the 95^th^-percentile of the Spearman *ρ*^2^ permutation null, each a measure of spike regularity (200 ms fano factor^46^, local LV^47^, and CV2^48^). *d*_2_ (*ρ*^2^ = 0.74) outperformed the best of these (Fano-200 ms, *ρ*^2^ = 0.34) by 40 points on this measure of rank-based predictive power. To understand how *d*_2_ relates to other features of neural activity, we computed each metric in closed form across the space of stable second-order (AR(2)) models and compared it to *d*_2_ in the same *ϕ*_1_–*ϕ*_2_ plane (Fig. 1D, Fig. S2). Within any local region of this space, nearly every standard descriptor of spiking covaries with *d*_2_. However, these relationships are often not monotonic, and don’t hold across the entire map (Fig. S2). Taken together, these data support the hypothesis that initial proximity to criticality determines the rate of subsequent learning in the brain. However, prey capture could appear purpose-built for confirming the critical brain hypothesis^10,49,50^. The task is complex, integrative, and ever changing, matching the defining features of critical dynamics themselves. This raises the question of whether criticality might not be predictive of the many more repetitive and low dimensional motor patterns that comprise much of daily behavior^13^. Accordingly, we examined learning in a second, six-day task in the same mice where mastery is defined by precision and stereotypy: crossing a 10-rung ladder.

### M1 proximity to criticality predicts learning of a stereotyped motor task: ladder crossing

Briefly, 18/18 mice spent six days living in a custom homecage whose two halves were separated, raised, and connected by a suspended ladder equipped with capacitive touch sensors (Fig. 2A). Ad lib food was available on one side, and water on the other. Animals engaged extensively with the ladder, crossing a mean of 649 times per 24 h (Minimum 60 crosses / day, maximum 1560 crosses / day, median 622 crosses / day). Two mice were excluded due to equipment malfunction, leaving *N* = 16 mice for all subsequent analysis. The capacitive touch time series made quantification and classification of crossing behaviors robust. Consistent with a lack of either prior ladder experience or a hard-wired ladder-walking circuit, early crossing attempts were hesitant, irregular, and slow. Eventually, all mice exhibited a set of readily distinguishable cross types, including stall-and-sniff, u-turns, walks, and highly stereotyped trots (Fig. 2B).

**Figure 2:**
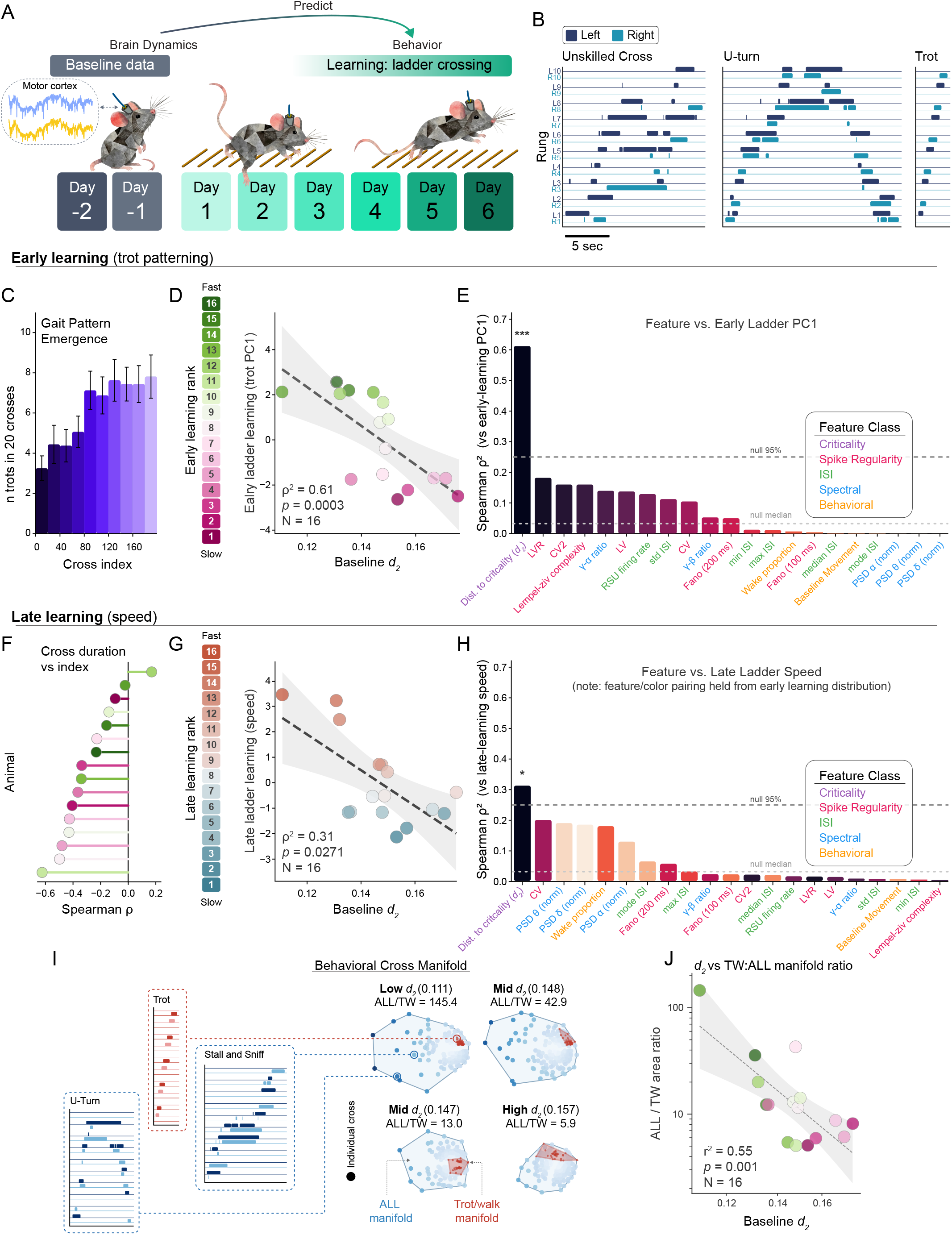
Proximity to criticality predicts ladder-cross learning rate. Across wild-type mice (*N* = 16), an individual’s baseline distance from criticality in M1 predicts how it discovers and then refines a stereotyped motor sequence: ladder crossing. **(A)** Experimental design: M1 single-unit ensembles were recorded across two baseline days and used to predict learning over a subsequent six-day ladder-crossing protocol. **(B)** Representative rung-contact time series (capacitive touch sensors; left rungs, navy blue; right rungs, teal) for three cross types: an early unskilled cross, a U-turn, and a stereotyped trot. **(C–E)** *Early learning (trot patterning).* **(C)** Gait-pattern emergence: the number of trots per 20 crosses rises with cumulative ladder experience in the first 200 crossings. **(D)** Baseline *d*_2_ predicts the rate of trot acquisition, quantified as a four-measure composite (trot PC1; *ρ*^2^ = 0.61, *p*_perm_ = 0.0003, *N* = 16; dashed line, fit; shaded band, 95% CI); individuals are colored by early-learning rank. **(E)** Twenty one baseline neural, spectral, and behavioral features were evaluated for the capacity to predict the future learning of trotting across the ladder. Only *d*_2_ cleared the 95^th^-percentile permutation null (dashed lines, null median and 95^th^ percentile; asterisks). **(F–H)** *Late learning (speed).* **(F)** Animals accelerate crossing speed with experience: across each animal’s first 500 purposeful (trot/walk) crossings, the within-animal Spearman correlation between on-ladder duration and crossing index is negative in 15 of 16 mice (median *ρ* = 0.34). **(G)** Baseline *d*_2_ predicts the rate of speed refinement, quantified as a three-measure cross-duration composite (*ρ*^2^ = 0.31, *p*_perm_ = 0.0271, *N* = 16); individuals are colored by late-learning rank. **(H)** Among the same features, *d*_2_ remains the only predictor above the 95^th^-percentile null; feature identity and color are held from panel E to illustrate that the ordering of metrics reshuffles between early and late learning. **(I–J)** *Behavioral exploration and consolidation.* **(I)** Ladder crosses embedded by multidimensional scaling (MDS) of their touch sequences; the convex hull of an animal’s first 200 crosses (ALL manifold, blue) contains the trot/walk subspace (red). Four animals spanning the *d*_2_ range show that animals closer to criticality (lower *d*_2_) pair a larger exploratory ALL manifold with a tightly consolidated trot/walk core (larger ALL/TW ratio). **(J)** The ALL/trot-walk area ratio is negatively correlated with baseline *d*_2_ (Pearson *r*^2^ = 0.55, *p* = 0.001, *N* = 16; log *y*-axis; power-law fit selected by BIC): animals closer to criticality combine the broadest exploration with the most consolidated purposeful core.

In the six days of ladder crossing, two categories of learning were readily discernible: an early phase in which mice learned to trot (pattern emergence and consolidation; Fig. 2C), and a late phase in which mice exhibited their fastest successful crosses (refinement and further improvement). Early learning was evident in the first 100 ladder interactions, and was well described by multiple measures, such as the number of trots in this set and the number of crosses to achieve the n^th^ trot. These measures were highly concordant (Kendall’s *W* = 0.914), and we calculated a composite as the first principal component across four measures to avoid parameter selection bias (Fig. S7A-C). Within the first ~100 ladder interactions, all 16 animals learned to successfully trot, but exhibited a spread in early learning rates (for example, a 29-fold range in n crossings to the 2^nd^ trot).

While ladder trotting is repetitive, low in variance, and largely fixed in its spatial and temporal aspects, we reasoned that the ability of a naive neurobiological network to explore a space of possible solutions for the behavior would nonetheless be limited by the initial proximity to criticality. To test this, we asked if baseline proximity to criticality was sufficient to forecast the rate of early ladder learning (recall that 8 animals went from baseline immediately to the ladder, while 10 first completed a 6 d prey-capture paradigm). Consistent with prey-capture learning, baseline proximity to criticality strongly predicted the rate at which each mouse learned to trot (*ρ*^2^ = 0.612, *p*_perm_ = 0.0003, *N* = 16; Fig. 2D). The effect was robust to leaving out any single animal (leave-one-out Spearman *ρ* ∈ [−0.882, −0.732]) and held for each early-learning metric individually (Fig. S7B,C). We again examined the predictive power of a panel of 20 additional baseline features, none of which reached the 95^th^-percentile of the permutation null (*ρ*^2^ = 0.250) (Fig. 2E). The closest feature, LVR (local variation of rate; *ρ*^2^ = 0.182), was not significant but is noteworthy as the closest measures to *d*_2_ in prey capture also quantified variance^51^.

In parallel with learning to trot, animals sped up their directed ladder traversals (trots and walks) over the first 500 crossings: on-ladder duration fell in 15 of 16 animals (median Spearman *ρ* = −0.34; sign test *p* = 2.6 × 10*^−^*^4^), the lone exception (*ρ* = +0.17) being among the quickest at the outset (Fig. 2C). Considered over thousands of crosses, animals approached their own speed asymptotes. In principle, this latter phase might be independent of proximity to criticality, as it does not require exploration of a solution space, only the rapid execution of a previously learned motor pattern. We tested this by evaluating how well baseline proximity to criticality predicted a three-metric composite of cross speed, considered from all crosses generated in the six day protocol (Fig. S7D-F; note that there was neither a correlation between total crosses and *d*_2_ nor the number of crosses required to reach top speeds). In contrast to our hypothesis, baseline proximity to criticality accounted for 31% of variance (*ρ*^2^ = 0.31) in late learning as indicated by fast crosses (*p*_perm_ = 0.027, *N* = 16; Fig. 2G), albeit with a lower prediction strength than early learning (late and early learning are correlated (*ρ* = +0.60*, p* = 0.014)). As in early learning, distance from criticality was the only baseline measure that predicted outcomes beyond chance (Fig. 2H). While below chance, the relative ordering of the 20 additional features was different from their rank in early learning. As with prey capture, asymptotic performance for both trot learning and cross speed refinement were not predicted by *d*_2_ (Fig. S6C,D). Taken together, these data suggest that a network’s basal proximity to criticality establishes learning rate not only for tasks that are variable, integrative, and evolving, but also for those characterized by stereotyped patterning and refinement^52,53^. In simple terms, a brain’s proximity to criticality appears to control both the ease with which solutions can be found as well as the pace at which they are refined on repetition.

Networks at criticality have the widest possible search space while retaining the ability to stabilize (i.e., exploit) favorable patterns^7,54^. Because the motor cortex directly contributes to and constrains an animal’s behavior, we reasoned that the same phenomena should be evident in the geometric structure of behavioral exploration during learning. To test this, we analyzed each animal’s first 200 ladder interactions and calculated the geometric distance between crosses (MDS^55–57^; Fig. 2I). The complete set of 200 crosses comprised an *All* manifold whose area indicates the total breadth of ladder behaviors. A subspace, the *Trot/Walk* manifold, captures the regularity of purposeful crossings. These manifolds are analogous to the explore—exploit balance described in other forms of learning^58,59^. M1 dynamics that are near criticality would enable a larger repertoire of neuronal patterns, which should in turn expand the behavioral repertoire and produce a broader *All* manifold. In alignment with this predict, we found that M1 baseline *d*_2_ was negatively correlated with the All manifold area (*ρ*^2^ = 0.32, *p_perm_* = 0.025; Fig. S8C); animals whose motor cortices were closer to criticality explored a larger range of ladder behaviors. Simultaneously, baseline *d*_2_ was *positively* correlated with trot/walk manifold area (*ρ*^2^ = 0.25, *p_perm_* = 0.047; Fig. S8D), suggesting that more critical animals identified and consolidated a solution set more rapidly; representative embeddings spanning the *d*_2_ range are shown in Fig. 2I. Considered together, these findings suggest that proximity to criticality offers a mechanism linking exploration and exploitation. Consistent with this, the ratio of All to Trot/Walk manifolds was negatively correlated with baseline *d*_2_ (Fig. 2J; Pearson *r*^2^ = 0.55, *p* = 0.001 on log–log axes, power-law fit selected by BIC because the All/Trot-Walk ratio is an unbounded, magnitude-meaningful quantity): animals closer to criticality paired the broadest exploratory repertoire with the most tightly consolidated purposeful core.

Taken together, the mouse data suggest that proximity to criticality is a stable feature of network dynamics that distinguishes animals and constrains subsequent learning. The two tasks were chosen to be maximally dissimilar, potentially indicating a general principle for how neural dynamics constrain learning. However, an equally plausible alternative explanation is that *d*_2_ captures a latent variable related to task engagement necessary for an animal to demonstrate improved performance but distinct from the mechanisms of learning.

### Proximity to criticality predicts plasticity of single neuron tuning

Ultimately, proximity to criticality should determine the ability of a complex network to generate the correlations that drive associative plasticity, whether in synthetic or biological systems. To directly test this, and sidestep confounding factors such as attention and motivation, we capitalized on the long-understood role of patterned visual experience in shaping isocortical responses^61^.

In ferret primary visual cortex (V1), single neuron direction selectivity emerges in an experience dependent, experimentally controllable fashion^62^ (Fig. 3A). According to our hypothesis, the magnitude of plasticity that stimuli can drive should be regulated by proximity to criticality. To test this, we combined previously published data^60^ (*N* = 5) with ten new recordings from ferret kits. To isolate stimulus-driven plasticity (eliminating variability in behavior and brain state), kits were lightly anesthetized while the activity of V1 neurons was recorded. Kits were then exposed to drifting gratings for multiple hours, after which increases in direction selectivity were robustly evident, albeit to varying degrees across animals (Fig. 3 B,C; slope = +0.0095 (1−DirCV) h*^−^*^1^, *p* = 3.0 × 10*^−^*^7^; linear mixed effects model: direction selectivity ∼ exposure time + (1 | animal), where animal is included as a random effect; estimated marginal means of 0.12 and 0.21 at 0 h and 9 h, respectively; *n* = 12 motion animals).

**Figure 3:**
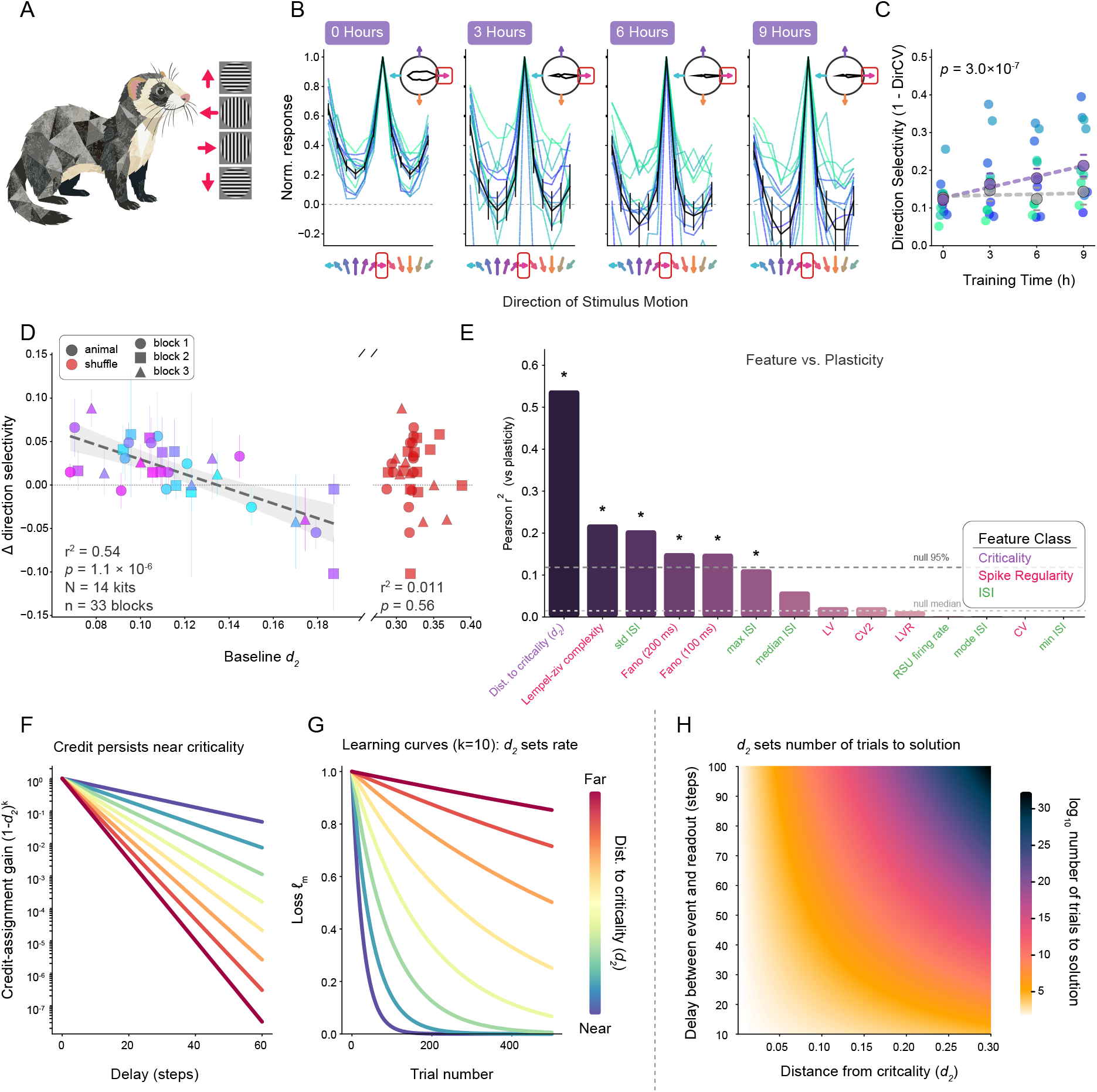
Proximity to criticality predicts developmental plasticity in ferret kits and defines learning rate in a synthetic learning model. **A-C**: Naive ferret kits (*N* = 15) exposed to moving stimuli **(A)** over hours **(B)** exhibited rapid increases in direction selectivity (individual lines represent animal-level means) **(C)** in ferret visual cortex^60^. **(D-E)**: Proximity to criticality predicts the degree of plasticity of V1 units per animal. Across a total of 33 blocks that met inclusion criteria, the degree of change in direction selectivity for units measured in that block was predicted by baseline *d*_2_ for V1 while no correlation was observed in the shuffle control **(D)**. Compared to 13 comparator features available in the ferret dataset, baseline *d*_2_ was the strongest predictor of the degree of plasticity in direction selectivity. **(F-H)**: A simple recurrent neural model with back-propagation reveals that proximity to criticality defines the speed for learning a task. The credit-assignment penalty associated with delay from input is shallow when *d*_2_ is low but steepens markedly as *d*_2_ increases **(F)**. For a fixed delay (k=10), the number of trials required to minimize loss to a minimal tolerance increases markedly with distance from criticality **(G)**. When the trials-to-solution surface is considered for a range of delays (k) and across a spectrum of *d*_2_ values, the number of trials to the solution increases super-exponentially **(H)**.

We calculated *d*_2_ from ensembles of V1 units prior to training blocks, allowing us to regress initial conditions against the degree of subsequent plastic change in the tuning of those same neurons. Notably, and consistent with M1 dynamics in freely behaving mice, baseline *d*_2_ in kit V1 exhibited a range of values across animals. Individual pre-stimulus-block measurements spanned 0.068 − 0.187; per-animal means 0.087 − 0.150, a 1.7-fold range. Thus, even in the context of uniform anesthetic application, animals populated a continuum of proximities to criticality that was wider than the spread of spike-shuffled control *d*_2_ (2.4-fold greater variance; Levene’s test, *p* = 0.037) and that served as the predictor axis for subsequent plasticity.

Directly contradicting our alternate hypothesis — that *d*_2_ might reflect some aspect of desire or engagement — pre-stimulus *d*_2_ in V1 was significantly negatively correlated with the pending change in direction selectivity (Fig. 3D; linear mixed-effects model, plasticity ∼ *d*_2_ + (1 | animal): *β* = −0.86, *p* = 1.1 × 10*^−^*^6^; *r*^2^ = 0.54 across *n* = 33 blocks from *N* = 14 kits whose data met inclusion criteria; exposure block did not modify the relationship). The effect held when collapsed to one independent point per animal (*N* = 14: Pearson *r* = −0.66, *r*^2^ = 0.43, *p* = 0.010). Conceptually, these results directly parallel the mouse ladder and prey-capture data; across species, tasks, and developmental stages, animals closer to criticality exhibited more experience-dependent change. Further, of the 20 additional baseline factors, 13 were relevant to the anesthetized kits (behavior and spectral features were excluded, as animals were not behaving and we only had spike times from previous work). Consistent with both mouse behavior paradigms, proximity to criticality far outperformed the second highest ranked feature (Fig. 3E). In this case, that was Lempel-Ziv complexity, which ranged from fourth to the least predictive feature across our other experiments. *d*_2_, in contrast, was the primary feature in every single context.

Taken together, our data are consistent with the hypothesis that proximity to criticality mechanistically determines plasticity rate. However, in complex, interacting systems, mechanism is challenging to define and address while avoiding mereological fallacy^63^; the *phenomenon* defines the level at which the mechanism operates^64,65^. For our purposes, the phenomenon is learning capacity, and the mechanism that explains it operates at the level of population dynamics: proximity to criticality appears to constitute the organized dynamical condition under which maximal information processing — and thus maximal plasticity — is possible. This level of organization is independent of lower details, thus the mechanism should hold equally in synthetic as in biological learning systems. Synthetic learning systems afford the opportunity to manipulate distance from criticality independently of the task and the learning rule.

#### Proximity to criticality constrains learning rate

In the present work, we use learning rate in the behavioral sense: the rate at which task performance improves with experience. This is different from the commonly known gradient descent step size *α*, which is a parameter controlling the size of each weight update. To ask how *d*_2_ could shape learning rate in a generic learning system, we turned to a minimal recurrent model for learning (Fig. 3F–H). Mathematically, even low-dimensional dynamical systems can exhibit criticality when approaching a bifurcation^31^. At a critical bifurcation, such systems exhibit critical slowing down, i.e., autocorrelation time diverges. In simple terms, the memory of past events persists longer the closer the system is to a bifurcation^66^. In a simple recurrent neural network trained by gradient descent, this same timescale sets the limit on temporal credit assignment: for an event *I* that happened *k* time steps in the past, the present error signal is governed by the product of recurrent dynamics, reducing to *µ^k^* (product of Jacobians), such that it vanishes (*µ <* 1) or explodes (*µ >* 1) unless *µ* is 1.0^67,68^. Over long timescales, then, we can reduce the RNN to its dominant real mode which gives a scalar recursion *h_t_* = *µ h_t−_*_1_ + *I_t_*. This is the definition of an AR(1) process for which the same distance-to-criticality we measure in neural data reduces to *d*_2_ = 1 − *µ* with *µ* ∈ (0, 1); the model’s distance from its bifurcation is measured directly by *d*_2_. For a linear readout learning a delayed association by gradient descent under squared-error loss, the number of trials needed to reach a fixed error tolerance *ɛ* (with asymptote *ℓ^∗^* = 0) is

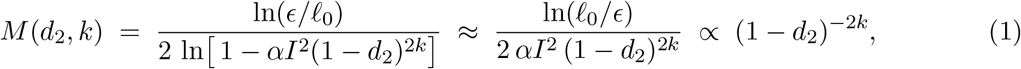

where *ℓ*_0_ is the initial loss, *α* the gradient descent step size, *I* the input drive at the first time step, and *k* the temporal horizon over which credit must be assigned (see Methods).

This one-parameter model reproduces the three principles that organize the animal data. First, it makes explicit the mechanism by which proximity to criticality matters: the credit-assignment gain (1 − *d*_2_)*^k^*— the fraction of an error signal that still reaches an event *k* steps in the past — decays geometrically with delay, with a decay constant fixed entirely by *d*_2_ (Fig. 3F). Near criticality the gain persists across long delays, so that temporally distant events remain visible to learning; away from criticality the gradient vanishes within a few steps, and the network is effectively blind to its own recent history. Second, feeding this surviving gradient into gradient descent determines learning *rate*, not asymptote, validating the principle that the speed of learning (but not its ultimate outcome) depends on *d*_2_. Every network can in principle reach zero loss, but the speed of that descent scales with (1 − *d*_2_)*^−^*^2*k*^: near-critical networks converge in tens of trials while more distant ones require hundreds to thousands, all settling onto the same floor (Fig. 3G). This is precisely the dissociation observed in the animals — baseline *d*_2_ forecasts how quickly a mouse improves and how much a kit’s tuning changes, but not the endpoint it eventually reaches. Third, because the cost scales as (1 − *d*_2_)*^−^*^2*k*^, the penalty for being off-critical compounds with task difficulty. The trials-to-solution surface (Fig. 3H) shows that for short credit horizons, slightly non-critical networks can learn in a feasible number of trials^13^, but as *k* grows the required trials rise exponentially with distance from criticality.

That a one-parameter, generic recurrent network reproduces the same *d*_2_–learning relationship seen across two mouse tasks and ferret visual plasticity indicates that the causal link between baseline proximity to criticality and the capacity for plastic change is a generic property of learning in recurrent systems operating near criticality.

### Proximity to criticality predicts general intelligence in humans

Much of human intelligence is necessarily acquired: one cannot speak a language without having been exposed to it and having undergone durable, experience-dependent change to retain it. Such intelligence is, by definition, a product of plasticity. This distinguishes it from capacities such as the stretch reflex, which is present in neonates and requires no experience to elicit^69^. Our animal and modeling data raise the possibility that, if high-resolution cortical electrophysiology were available in relevant human circuitry, proximity to criticality might provide a mechanistic explanation of durable, inter-individual variation in cognitive performance. Because human validation of our hypothesis requires both high-resolution neural data and normed cognitive assessment, analysis necessitates a clinical cohort. With this in mind, we examined a stereoelectrode-based electrophysiology dataset collected from 35 patients undergoing chronic monitoring in a children’s epilepsy clinic (Fig. 4A). As *d*_2_ can be measured in as little as 1,000 samples^40^, we curated a relatively small sample (∼ 60 min) of artifact free data from a frontal isocortical probe in each patient during waking. Note that, proximity to criticality in EEG is indistinguishable between epilepsy patients and healthy controls when taken from inter-ictal periods removed from seizure activity^70^. In our dataset, each patient’s full scale intelligence quotient (IQ) was assessed by clinicians. While IQ collapses heterogeneous abilities into a single score and has a history of misuse^71^, it reliably and predictively indexes a durable, high-level capacity for adaptive cognition: full-scale scores are among the most stable and reproducible measures in all of psychology, with high test–retest reliability, rank-order stability that persists across decades^72^, and predictive validity for educational, occupational, and health outcomes that exceeds that of any comparable single variable^71,72^.

**Figure 4:**
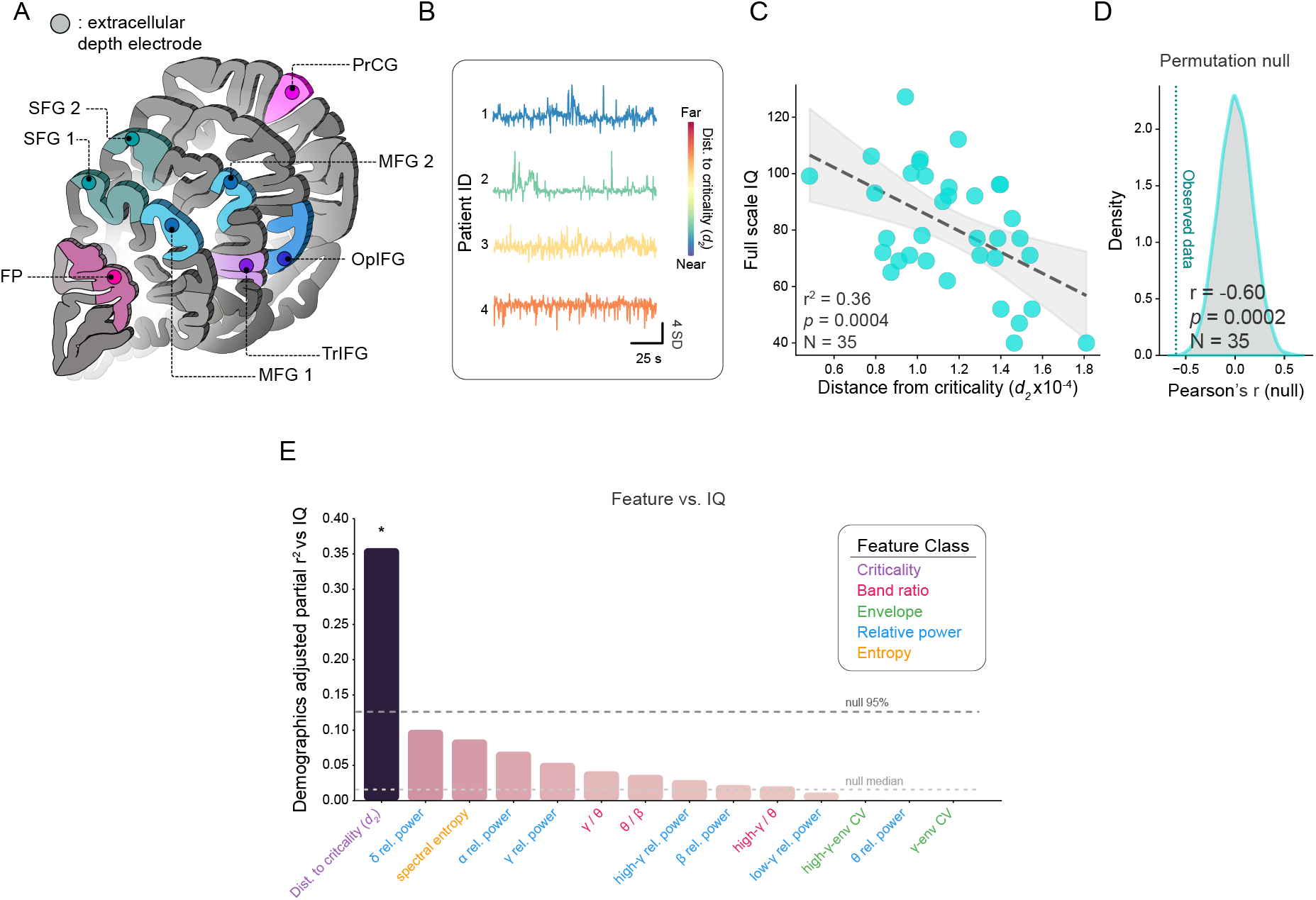
Proximity to criticality predicts general intelligence in human patients. **A**: Frontal isocortical entry sites of the analyzed stereoelectrode rods, shown on representative coronal plates from the Allen Human Brain Atlas^73,74^ (Allen Institute for Brain Science; human.brain-map.org). Each marker denotes the superficial cortical entry of one patient’s analyzed rod and indicates regional coverage rather than an exact contact coordinate. Note that one brain region (PrCG, *N* = 2 patients) is not represented as this region is rostral to the first available Allen Human Brain Atlas plate. For another patient (*N* = 1) the region is unassigned, though confirmed to be in the frontal cortex **(B)**: Exemplar enveloped stereo EEG recordings show multiple timescales that vary with proximity to criticality. **(C-D)**: Isocortical proximity to criticality predicts IQ in a clinical epilepsy population (*N* = 35) **(C)** and a permutation test confirms that this relationship could not have arisen by chance **(D)**. **(E)**: Comparison of *d*_2_ against comparator features reveals that proximity to criticality is the only neural feature able to predict IQ in our dataset.

Importantly, IQ tests probe the ability to solve novel problems, not previously acquired knowledge or skills. Thus, they aim to quantify the capacity to acquire and integrate, which is, conceptually, the plastic operation our data suggest arises from proximity to criticality. To test this directly, we computed *d*_2_ from frontal stereo EEG. Because clinical stereo EEG resolves mesoscale local field potentials rather than single units, we fit the autoregressive model directly to continuous data; the high-gamma envelope serves as a proxy for local multi-unit bursting^75,76^, and its long-range temporal correlations are preserved beyond the recording hardware’s high-pass cutoff (Fig. 4B). In the patient cohort, frontal cortical proximity to criticality correlated with full-scale IQ: lower *d*_2_ was associated with higher IQ after adjusting for demographic covariates (age, sex, race/ethnicity, and handedness; partial *r* = −0.60, *r*^2^ = 0.36, *p_perm_* = 3.8 × 10*^−^*^4^, *N* = 35; Fig. 4C,D), an effect that was already significant at the zero-order level (Pearson *r* = −0.51, *r* = 0.26, *p* = 1.7 × 10^-3^ ). In contrast, in our cohort of epileptic patients, full-scale IQ did not vary significantly with seizure burden (Spearman *ρ* = −0.12, *p* = 0.49, *N* = 35). Similarly, although in this cohort IQ increased with age across the 2–22 y range (*r* = +0.39), *d*_2_ was uncorrelated with age (*r* = −0.17, *p* = 0.34), and differed neither by sex (point-biserial *r* = +0.01, *p* = 0.94; Cohen’s *d* = 0.03), nor race/ethnicity (point-biserial *r* = +0.01, *p* = 0.97; Mann-Whitney *p* = 1.00).

As in the animal experiments, we asked whether this prediction generalized to other features of the signal. Because sEEG resolves mesoscale field potentials rather than spikes - and because absolute signal amplitude is not comparable across electrodes - we competed *d*_2_ against 13 amplitude-invariant spectral features. Of the set, only *d*_2_ predicted IQ above chance (next-best feature *r*^2^ = 0.10, *p*_perm_ = 0.07; Fig. 4E), mirroring its dominance over the metrics evaluated in mice and ferrets.

Previous work has shown an association between proximity to criticality and general intelligence in adults and children, based on functional imaging or surface EEG-based measures^77–79^. Our results extend this literature by using direct intracranial recordings alongside a time-domain criticality measure, and we found a strong predictive relationship. When considered alongside the mouse, ferret, and model data, our findings argue that isocortical proximity to criticality constrains a common capacity for experience-dependent change, conserved from mouse to human.

## Discussion

The brain is the source of the adaptive, complex behavior required to thrive in an unpredictable world. Because behavior is the substrate for evolutionary selection^80^, animals that adapt more quickly and effectively should out-compete others. As a result, brains, regardless of anatomical specifics, might be expected to ultimately converge on the computational principles that maximize adaptability^10^. Through this lens, due to its direct, *a priori* explanatory power regarding behavior and function, criticality has been suggested as a plausible unifying feature of brain function across species and an endpoint of homeostatic plasticity^22,81^. Consistent with this, proximity to criticality degrades in pathologic states that compromise adaptability^82^. What has remained untested, however, is the central prediction that this endpoint constrains adaptive plasticity in the brain. Here, we provide multiple lines of evidence supporting that conclusion. We find that, just as with other homeostatic set-points in physiology^41,42,83^, in vivo brain networks exhibit a range of near-critical set-points that are stably maintained at the level of the individual. Animal-level variation within this range appears to consequentially determine the rate at which an individual can drive plastic change, such as in learning, in the future. In complex systems not amenable to interventionist tests, such as economies^84^ or ecosystems^85^, consistent predictive power is a test of causality^86^. To go beyond this, we establish causality using a synthetic learning model. Finally, we present evidence for the existence of a similar relationship in the adaptability of the human brain.

Our findings also speak to the function of sleep. Sleep is required for adaptation across the animal kingdom, from respiratory plasticity^87^ to complex cognition^88^ — a universality that is hard to explain if sleep services a catalog of circuit-specific needs. Occam’s razor favors a single shared variable, necessary for adaptive computation and restored by sleep; proximity to criticality fits, being homeostatically maintained^22^, eroded across waking and renewed by sleep^23,89^, and progressively, sleep-dependently lost in neurodegeneration^82^. Tellingly, although innumerable features differ between sleep and wake, it is distance from criticality — not slow-wave energy or any other measure— that predicts impending state transitions^23^. Our results demonstrate that the same set-point governs the rate of future plastic change, the very capacity that fails without sleep.

The same logic links dynamics and learning. Prior work has tied spectral features^90,91^, neural complexity^92^, and spike-time irregularity^93^ to learning, plasticity, or cognitive ability. Our analytical and empirical results show these are subsumed by proximity to criticality. Because *d*_2_ is a geometric property of the population’s dynamics, many such features must covary with it. The covariance of *d*_2_ and other features tends to be local, as the relationships shift across the space (Fig. S2), such that none reconstructs *d*_2_ and *d*_2_ cannot be trivially built from any of them. However, *d*_2_ does not exist in a vacuum: autocorrelation time offers a familiar though not independent bridge to distance from criticality^94^. Autocorrelation time and susceptibility, like long-range temporal correlations^34,95^ and the branching ratio^96,97^, are signatures of an RG fixed point, each diverging or becoming singular as the system nears one (critical slowing down). Each offers an indirect readout of proximity to criticality, whereas *d*_2_ measures that distance directly in the space of AR models. As a confirmation, a panel of autocorrelation-based statistics (integral and timescale, AR-dominant timescale, and susceptibility) also predicted learning across all three tasks, and theirv theoretical dependence on the AR coefficients tracks the critical boundary but less directly and less uniformly than *d*_2_ (Fig. S9). Because they are indirect, their contribution was variable within and across tasks: *d*_2_ remained the strongest predictor for both trot-emergence and prey-capture learning, and was surpassed only in late-phase ladder refinement (speed) — the setting in which we had predicted the smallest role for criticality and observed the weakest *d*_2_ effect.

The deepest implication of our results is not specific to a task, region, or species, but the relationship between dynamical organization and computation itself. It is increasingly clear that a system’s capacity to learn is constrained by mathematical principles that are largely indifferent to its physical substrate; learning arises in cephalopods^98^ as it does in LLMs. The same distance-to-criticality that governs credit assignment in our recurrent model appears to govern learning rate in three unique mammalian brains. This advances the hypothesis^11,50^ that proximity to criticality is not an abstract correlate of cognition but constitutes the dynamical regime that accounts for the adaptive computation underlying much of behavior and brain function. If so, the principle should extend to any aspect of cognition that depends on plastic change. We emphasize, however, the boundaries of what our data establish. The relationship is predictive across species and causal within a model; a direct demonstration that manipulating proximity to criticality alters learning rate in the brain remains untested. Further, the human association is cross-sectional, and whether the same principle holds in nervous systems that evolved independently of the mammalian cortex is unknown, although prior work describes near-critical dynamics in, e.g., leeches, fish, and turtles^10^.

Proximity to criticality thus offers a mechanistic account of variability in learning rate at the level of population dynamics^65^, and represents a theory-derived prediction^1,4,99^ borne out across species, regions, and paradigms. It does not specify the lower-level machinery that establishes the set-point — synaptic homeostasis^100^, cell-non-autonomous network governance^101^, and the sleep-dependent restoration described above^23^ are candidates. However, theory and accumulating empirical evidence suggest that diverse underlying machinery establishes a unifying, emergent computational regime necessary for experience-dependent plastic change. From this perspective, each cellular and molecular component bears on learning only insofar as it moves the brain toward or away from criticality, or establishes the capacity for plastic modulation of connection weights (e.g., input-specific associative plasticity). In simple terms, proximity to criticality may reflect the speed limit of adaptation in complex learning systems, whether biological or artificial.

## Supporting information

Early ladder crossing

Trot crossing, viewed from 1 angle

Trot crossing, viewed from the other angle

Trot crossing at 0.25x speed

U-turn crossing

Vertical exploration crossing

## Methods

### Mice

All procedures were performed in accordance with protocols approved by the Washington University in Saint Louis Institutional Animal Care and Use Committee, following NIH guidelines for the Care and Use of Laboratory Animals. In total, 12 female and 6 male adult wild-type C57BL/6 mice were used (Jackson Labs, strain #000664). Mice were postnatal day 65 (P65) to P123 at the onset of baseline recording (mean = 80.0 d, s.d. = 15.9 d). All mice were housed in an enriched environment on a 12:12 h light/dark cycle.

### Surgery

Custom-built multielectrode arrays were assembled and implanted as described previously^23,39,82^. Prior to surgery, mice were administered buprenorphine SR (0.1 mg kg*^−^*^1^), dexamethasone (0.5 mg kg*^−^*^1^), and meloxicam (5 mg kg*^−^*^1^). Mice were anesthetized with isoflurane (3% for induction, 1–2% for maintenance, in air) and head-fixed in a robotic stereotaxic instrument. The fur, skin, and periosteum over the dorsal skull were removed, and a 1–1.5 mm craniotomy was cut over the target region. Custom-built, 64-channel, tetrode-based arrays were lowered into each target at 5 mm min*^−^*^1^ using a robot and vacuum-based probe holder. All 18 mice received a 64-channel array in the right M1 caudal forelimb area (coordinates, AP/ML/DV relative to bregma and dura: 0.25*/*1.25*/*0.79 mm). All animals were implanted in the right hemisphere so that hemisphere was held constant across the cohort, removing inter-hemispheric variability as a confound in between-animal comparisons of *d*_2_; the right hemisphere was chosen arbitrarily. In 5 of the 18 mice, an additional, second 64-channel array was implanted in right primary visual cortex (V1; coordinates^39^ −3.20*/*2.20*/*1.00 mm), targeted as in Parks et al., yielding 128 channels across the two regions. Unless otherwise noted, analyses reported here use the M1 ensembles. A bridged ground/reference wire was connected to a skull screw over the contralateral hemisphere; the array was connected to headstage electronics (White Matter LLC) via custom PCB-EIBs. Electronics were enclosed in a 3D-printed case, and secured with dental cement. The entire assembly weighed *<* 3 *g*. Mice received meloxicam (5 mg kg*^−^*^1^) and dexamethasone (0.5 mg kg*^−^*^1^) for 3 d post-operatively and recovered for a minimum of 7 d before recording began.

### Recording

Recordings were performed continuously in freely behaving mice across three environments: a standard home cage during the baseline period, a circular prey-capture arena, and a custom ladder cage (both described below). Throughout recording, mice were connected via a custom cable containing an in-line commutator (slip ring) that permitted unrestricted movement. Neural signals were amplified, digitized (14-bit), and acquired at 25 kHz on the eCube acquisition system (White Matter LLC). Behavioral video was captured at 15 or 30 frames s*^−^*^1^ and temporally aligned to the neural data. For each animal, a continuous 13–14 d recording (312–336 h) comprised a 24–48 h baseline of spontaneous home-cage behavior followed by two 6-day motor-learning tasks; task order (prey capture first vs. ladder first) was randomized across animals.

### Prey-capture task

Prey were young-adult Turkestan red-runner cockroaches (*Shelfordella lateralis*; Caribbean Mealworms) weighing 0.05–0.20 g (mean = 0.145 g, s.d. = 0.041 g). For prey-capture experiments, mice were housed individually in circular hunting arenas constructed from 19 L (5-gallon) buckets, which served as home cages for the 6 d protocol. On Day 1 (pre-trial), a single cockroach was introduced at 17:00 and left overnight; any cockroach not captured was removed the following morning at 09:00. On Day 2, food (but not water) was removed at 17:00, beginning a 16 h food-deprivation period. Beginning at 09:00 on Day 3, cockroaches were presented one per trial, up to six trials per day. Each trial lasted *<* 30 min: if a mouse did not engage or failed to hunt successfully, the cockroach was replaced. If a mouse captured its prey rapidly, a 10 min delay was introduced before the next cockroach was presented. This 16 h cycle of food deprivation and hunting was repeated on four consecutive days; each mouse encountered 25 cockroaches in total (4 days × 6 trials, plus the pre-trial cockroach). Mice had *ad libitum* access to food from Day 1 through the evening of Day 2; water was available *ad libitum* at all times except during the four hunting sessions.

### Automated ladder-rung walking task

To embed a structured motor challenge within the home cage, we developed a self-paced ladder paradigm in which a horizontal ladder separates the food and water sources, so that the mouse must traverse it to feed and drink (Fig. 2B). The ladder was made from laser-cut acrylic and T-slotted aluminum extrusion (80/20). Mice were kept on a 12:12 h light/dark cycle (visible lights on at 07:00, off at 19:00), with infrared lights on continuously so cameras could image animals throughout the dark phase. Each ladder cross constituted a single “trial,” yielding a naturally repeating trial structure while preserving spontaneous, ethological behavior. Over days, mice generated thousands of crosses spanning the full course of learning, from initial acquisition through a stabilized motor solution.

Behavior was captured from both sides of the ladder by two Raspberry Pi High Quality (HQ) cameras (12 MP) recording at 60 fps, each driven by a dedicated Raspberry Pi 4B. Rung contacts were sensed by two MPR121 12-channel capacitive touch controllers (breakout boards, Adafruit; NXP MPR121 IC) sampled at 1 kHz and read by a single Raspberry Pi 4B. We used 20 channels to encode 10 left–right rung pairs. To permit simultaneous neural recording, the capacitive sensors were electrically isolated from the controlling Raspberry Pi with bidirectional, I^2^C-compatible digital isolators (ISO1640, Texas Instruments), preventing bus noise from contaminating the neural recording. All data streams—video-based limb kinematics and rung-contact times—were synchronized to the electrophysiological recording. Inspired by the Precision Time Protocol^102^, a Raspberry Pi Pico broadcast a synchronization pulse to all recording devices every 10 s, and the streams were aligned post hoc to this shared reference clock.

### Spike sorting and unit classification

Only baseline-period neural data were analyzed in this study. Broadband data were band-pass filtered between 500 and 7,500 Hz. Candidate spikes were detected by a threshold between 4.5 and 6 standard deviations below the mean of the filtered signal; for each animal, the threshold yielding the greatest number of single units was selected. Spikes were sorted with a modified implementation of MountainSort5^103^ and SpikeInterface^104^ in non-overlapping 4 hour clustering blocks. Clusters were classified by an XGBoost model using features including waveform shape, waveform consistency, inter-spike-interval distribution, and multi-channel spatial spread^105^. Cluster quality was confirmed manually by trained scorers blinded to behavioral outcome. Each unit was assigned to one of four quality tiers; all analyses were restricted to well-isolated single units (Quality 1 and 2), excluding multi-unit (Quality 3) and noise-contaminated (Quality 4) clusters. Single units were further classified as regular-spiking (RSU) or fast-spiking on the basis of spike waveform^106^. All analyses reported here include only RSUs, and blocks with *n* ≤ 15 RSUs were excluded from analysis owing to inaccurate measurement of *d*_2_ from such a limited set. Across the 18 mice, 6,326 RSUs were recorded (mean = 34 ± 13.9 per animal; range 15–71).

### Arousal-state classification

Arousal state was manually scored from a combination of spectral and movement features^23,39^. For each recording, three M1 channels with clean local field potential (LFP) were downsampled to 500 Hz, and LFP power in the 0.1–60 Hz band was rendered as hourly spectrograms. Synchronized behavioral video was processed with DeepLabCut^107^ for pose estimation, and the resulting movement traces were aligned to the LFP. States were assigned in 4 s epochs consistent with well-established parameters^37,108^: high delta-band power with low movement was scored as NREM sleep; low delta-band power with elevated theta-band power and no movement was scored as REM sleep; and elevated theta-band power accompanied by movement was scored as wake. In total, 728 h of baseline recording across the 18 mice were scored manually. Unless otherwise noted, all neural features were computed exclusively during waking epochs.

### Ferrets

#### Subjects and general design

Ferret data were collected in the laboratory of S.D.V.H. at Brandeis University under protocols approved by the Brandeis University Institutional Animal Care and Use Committee, following NIH guidelines. The dataset comprised 15 female ferret kits (*Mustela putorius furo*), aged postnatal day 31–34 (around the time of natural eye opening, when direction selectivity begins to emerge), studied in terminal, anesthetized electrophysiology experiments.

Twelve kits received visual training with drifting gratings and three received a static gray-screen control. Of the 15, five were reported previously^60^ and the remaining ten were newly recorded. Female animals were used exclusively because animals were co-housed with sexually mature females and co-housing with males induces stress^60^.

#### Surgery and anesthesia

Kits were initially sedated with ketamine and given atropine and dexamethasone to reduce secretions and inflammation. Anesthesia was induced with isoflurane in a nitrous-oxide/oxygen mixture delivered by mask, and a tracheostomy was performed for mechanical ventilation (1.5–3% isoflurane in 2:1 nitrous oxide:oxygen). An intraperitoneal cannula was placed for delivery of Ringer’s solution and the neuromuscular blocker gallamine triethiodide, which suppressed spontaneous eye movements during recording. The electrocardiogram was monitored continuously to titrate anesthetic depth, and body temperature was held at 37 *^◦^*C. A 4 × 4 mm craniotomy was made over primary visual cortex (V1) in the right hemisphere and the dura was resected.

#### Electrophysiology

Multiunit activity was recorded across all cortical layers of V1 with a 32-channel silicon probe (NeuroNexus, A1x32-Poly2-10mm-50-177) inserted approximately perpendicular to the cortical surface (900–1100 *µ*m depth), with agarose applied to stabilize the brain. Signals were acquired on one of two systems (11 animals via a Multichannel Systems preamplifier/amplifier with LabVIEW and a National Instruments 6071e board; 4 animals via an Intan RHD2132 headstage). For both systems, spike waveforms were detected at a threshold of 5 standard deviations, and all threshold crossings on a channel were pooled as a multiunit cluster, because activity attributable to isolated single units was rarely resolvable with these electrodes^60^.

#### Visual stimulation and direction-selectivity assessment

Visual stimuli were generated in MATLAB with the Psychophysics Toolbox and displayed on a Sony GDM-520 monitor 45 cm from the animal. Stimuli were full-field, high-contrast drifting sinusoidal gratings presented in 12 directions (30*^◦^* increments), with a spatial frequency of 0.08 cyc deg*^−^*^1^. The experiment alternated assessment and training blocks: direction tuning was assessed with gratings at 25 deg s*^−^*^1^ before training (0 h) and after each of three 3 h training blocks (3, 6, and 9 h), during which the animal viewed drifting gratings (motion condition) or a static gray screen (control). At each assessment, direction selectivity per recording site was quantified as 1 DCV, where DCV is the directional circular variance^109,110^. Sites were included only if they (i) showed significant response variation across stimuli by ANOVA, (ii) responded at ≥ 1 Hz to 25 deg s*^−^*^1^ gratings, and (iii) had a preferred orientation within 30*^◦^* of the optimal orientation. The whole-animal, per-block plasticity used below is the change in the cell-averaged 1 − DCV across an exposure block.

### Human stereo-EEG

#### Patients

We performed a retrospective, observational study of children and young adults, ages 2–25 y, who underwent clinically indicated stereotactic EEG (sEEG) monitoring to identify a seizure focus between February 2016 and July 2023 in the Epilepsy Monitoring Unit. The study was approved by the Washington University in St. Louis Institutional Review Board (Protocol 202412145); consent was waived as the study used observational, retrospective data. Electrodes were standard, clinically approved depth electrodes (800 *µ*m diameter, 2 mm contact length, 3.5 mm inter-contact spacing; PMT, Chanhassen, MN). Implantation was planned per patient according to the primary hypothesis for seizure localization, with adjacent electrodes demarcating the seizure-onset zone and additional electrodes placed to identify eloquent cortex for language and motor mapping.

From a total cohort of *N* = 54 patients recorded, *N* = 35 met inclusion criteria of (1) a valid full-scale intelligence quotient (IQ) and (2) artifact-free frontal recordings during waking. These 35 patients (ages ∼ 2–22 y) are summarized in detail in Table 1. Full-scale IQ was determined for each patient by clinical neuropsychological assessment (WISC-V or WAIS-IV, age-appropriate version; administered by a licensed clinical neuropsychologist) as part of standard presurgical epilepsy evaluation.

**Table 1:** Analyzed patient demographics. Summary of the analyzed cohort (*N* = 35), each contributing a single frontal isocortical contact. Continuous variables are reported as median [IQR] with range; categorical variables as count (%). Seizure burden reflects the clinical cvEEG estimate (approximate seizures per day). ASD, autism spectrum disorder; TSC, tuberous sclerosis complex; FSIQ, full-scale IQ; IQR, interquartile range.

| Characteristic | Value |
| --- | --- |
| *Age (years)* |  |
| Median [IQR] | 14 [10–18] |
| Range | 2–22 |
| *Sex* |  |
| Male | 20 (57%) |
| Female | 15 (43%) |
| *Handedness* |  |
| Right | 26 (74%) |
| Left | 9 (26%) |
| *Ethnicity* |  |
| White, non-Hispanic | 31 (89%) |
| Black/African American, non-Hispanic | 3 (9%) |
| Asian | 1 (3%) |
| *Additional Diagnoses* |  |
| ASD | 7 (20%) |
| TSC | 6 (17%) |
| *Prior lesion resection* | 5 (14%) |
| *Seizure burden (approx. seizures/day)* |  |
| Median [IQR] | 1.6 [0.5–3.0] |
| Range | 0–20 |
| *FSIQ* |  |
| Median [IQR] | 77 [70–96] |
| Range | 40–127 |

#### Stereo EEG acquisition

Stereo-EEG uses intracranial *depth* electrodes (not scalp or subdural surface electrodes); each rod carries multiple contacts sampling local field potentials along its trajectory, typically on the order of 150 contacts per patient. For analysis we analyzed a single superficial electrode from each patient in a non–seizure-onset region with similar anatomic positioning across subjects (e.g., motor, frontal, or cingulate cortex). Anatomic regions were identified using CT, MRI, and Curry 3D modeling software (Compumedics, Charlotte, NC), and contact locations were reconstructed from clinically acquired imaging. EEG was acquired on a clinical Nihon Kohden system (Tokyo, Japan) using a non–DC-coupled amplifier with 250 channels, at a native sampling rate of 1,000 Hz (studies from 2016–2017 used 500 Hz); native rate varied by patient and was standardized to 500 Hz in analysis (see below). Anti-seizure medications were gradually weaned during EEG recording period. When possible, only epochs free of seizure activity were used. For some patient’s with abundant seizures, epochs with the lowest seizure burden were analyzed. In all cases, seizure activity was verified by a clinical epileptologist. Consistent with prior work, inter-ictal dynamics in epilepsy patients show normal signatures of criticality^70^. Data were deidentified and exported in European Data Format (EDF); clinical and demographic data were maintained by the clinical study team.

#### Contact selection and preprocessing

For each patient, a single frontal isocortical contact was analyzed, and approximately 60 min of artifact-free waking data were used. Across the cohort, analyzed contacts were distributed over frontal isocortical gyri of both hemispheres (per-patient localization in Table 2; entry sites shown on Allen Human Brain Atlas coronal plates in Fig. 4A): middle frontal gyrus (*N* = 12), inferior frontal gyrus pars opercularis (*N* = 12), superior frontal gyrus (*N* = 4), inferior frontal gyrus pars orbitalis (*N* = 2) and pars triangularis (*N* = 1), precentral gyrus (*N* = 2), and frontal pole (*N* = 1); one patient carried a bilateral implant that could not be assigned to a single gyrus.

**Table 2:** Localization of the analyzed frontal sEEG contact, per patient. Each patient contributed a single frontal isocortical contact (*N* = 35). Hemisphere: L, left; R, right; U, undetermined. Region abbreviations: MFG, middle frontal gyrus; SFG, superior frontal gyrus; OpIFG, inferior frontal gyrus (pars opercularis); TrIFG, inferior frontal gyrus (pars triangularis); OrIFG, inferior frontal gyrus (pars orbitalis); PrCG, precentral gyrus; FP, frontal pole. One patient carried a bilateral implant that could not be assigned to a single gyrus. Patient identifiers are the study’s internal codes.

| Patient | Hem. | Region | Patient | Hem. | Region |
| --- | --- | --- | --- | --- | --- |
| 4C | L | MFG | 3C | R | OpIFG |
| 17C | R | MFG | 6C | R | OpIFG |
| 20C | L | MFG | 14C | R | OpIFG |
| 34C | L | MFG | 16C | L | OpIFG |
| 40C | L | MFG | 21C | L | OpIFG |
| 41C | L | MFG | 26C | L | OpIFG |
| 42C | L | MFG | 30C | L | OpIFG |
| 49C | R | MFG | 38C | R | OpIFG |
| 52C | R | MFG | 43C | R | OpIFG |
| 55C | L | MFG | 48C | L | OpIFG |
| 56C | R | MFG | 59C | L | OpIFG |
| 57C | L | MFG | 60C | L | OpIFG |
| 36C | L | SFG | 45C | L | TrIFG |
| 50C | L | SFG | 39C | R | OrIFG |
| 53C | R | SFG | 47C | R | OrIFG |
| 58C | L | SFG | 31C | R | PrCG |
|  |  |  | 46C | L | PrCG |
|  |  |  | 23C | R | FP |
|  |  |  | 8C | U | bilateral (unassigned) |

Signals were resampled to 500 Hz (polyphase resampling), notch-filtered at 60, 95, and 120 Hz (IIR notch, *Q* = 30), and high-pass filtered at 0.1 Hz (4th-order Butterworth, zero-phase). Transient artifacts were removed within each analysis window by flagging samples whose amplitude exceeded the window median by 3 robust standard deviations (1.4826× MAD), padding flagged samples by 50 ms, and linearly interpolating gaps up to 1 s; windows in which more than 10% of samples were interpolated, or which contained any longer gap, were discarded. Cleaned data were segmented into non-overlapping 180 s windows for *d*_2_ estimation (below).

### Proximity to criticality (***d*_2_**): spike-based ensembles (mice and ferrets)

#### Definition

Proximity to criticality was quantified as *d*_2_, a distance-to-criticality measure derived from a renormalization-group^111^ treatment of univariate time series^40^. A population time series was fit with an order-*p* autoregressive (AR) model, yielding coefficients *ϕ*_1_*, …, ϕ_p_* that specify a point in the *p*-dimensional space of AR(*p*) models. The set of perfectly critical models—those generating autocorrelations across all scales—forms a hyperplan 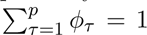 in this space, and *d*_2_ is the Euclidean distance from the fitted point to that hyperplane,

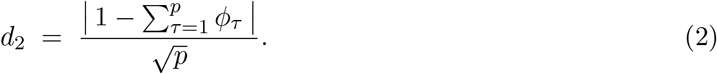

Throughout, *p* = 10. Smaller *d*_2_ indicates greater proximity to criticality; white noise (all *ϕ_τ_* = 0) gives a reference for ‘far from criticality’ 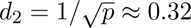 for *p* = 10. The same definition is used for mice, ferrets, and humans; only the substrate and the estimation of the AR coefficients differ.

#### Mouse *d*_2_

Only regular-spiking, Quality-1–2 single units were included. Each baseline recording was divided into non-overlapping 5 min windows, and only windows in which the animal was awake for *>* 90% of the window were analyzed. Within each window, ensemble spike counts were summed into 15 ms bins to form a population spike count time series. A tenth-order AR model was fit by the Yule–Walker method with maximum-likelihood estimation (statsmodels), and *d*_2_ was computed from the resulting coefficients by Eq. (2). Unless otherwise noted, each animal’s baseline *d*_2_ is the mean across its wake windows.

#### Ferret *d*_2_

For ferrets, *d*_2_ was computed per animal and exposure block from the pooled multiunit spike train, using the same distance-to-criticality definition (Eq.2) as for mice. Because ferret firing rates varied widely across recordings, the bin width DT was set per recording as a fixed multiple of the mean inter-spike interval, 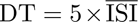. Analysis was restricted to “valid” windows of sustained activity, i.e., periods with ≥ 1 spike per bin sustained for ≥ 3 s—because including inter-stimulus and blank periods allows stimulus structure to swamp the signal. The autocorrelation function was accumulated within valid windows (averaged across all valid windows, with no cross-window lag pairs), and the AR(10) coefficients were obtained from this windowed autocorrelation function by Yule–Walker method; *d*_2_ then followed from Eq. (2). This estimator is algebraically identical to the mouse definition. Because the multi-hour anesthetized preparation can drift in depth, blocks were screened for anesthetic plane by two criteria: 1) the spontaneous/evoked activity ratio had to be below 0.7 to avoid periods approaching emergence from anesthesia, and 2) and the population spike rate had to exceed 13 spikes *s^−^*^1^, which excludes blocks suppressed under an overly deep plane and ensures sufficient spiking for a stable AR fit. Gray-screen control animals, which have no evoked response against which to define the ratio, were included under the rate criterion alone (see Fig. 3 analysis, below).

#### Spike-time shuffle control

To establish a white-noise upper bound on *d*_2_ empirically, we computed a shuffled control 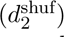 in which each neuron’s spike count within a window was preserved while its spike times were redrawn independently and uniformly at random across the window. Shuffled single-neuron trains were recombined into a population time series and *d*_2_ was recomputed as for intact data. This preserves each neuron’s firing rate while destroying temporal correlations across the population, driving 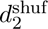 toward the analytic white-noise value 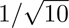 (mice, Fig. 1C; ferrets, Fig. 3D).

#### Proximity to criticality (***d*_2_**): field potentials (humans)

Clinical stereo EEG resolves mesoscale LFP rather than single-unit spikes. We therefore fit the AR model to a continuous field-derived signal: the amplitude envelope of the high-gamma band, which serves as a proxy for local multi-unit bursting^75,76^ and whose long-range temporal correlations are preserved above the recording hardware’s high-pass cutoff. Within each preprocessed 180 s window (above), a single frontal contact was band-pass filtered to the gamma band (70–250 Hz; 4th-order Butterworth, upper edge capped just below the 250 Hz Nyquist), and the analytic amplitude envelope was obtained by Hilbert transform and smoothed with a 100 ms Gaussian kernel. The envelope was *z*-scored (rendering *d*_2_ invariant to absolute signal amplitude) and fit with an order-10 AR model by Yule–Walker maximum-likelihood estimation. *d*_2_ was computed by Eq. (2). Each patient’s *d*_2_ is the mean across that patient’s 180 s windows; a 16-way aggregator sweep confirmed the mean is optimal for the effect and that the result is robust to this choice.

#### Baseline ***d*_2_** setpoint and its stability (Fig. 1E, S3)

Because *d*_2_ varies systematically by arousal state^40^, baseline analyses focused on wake to estimate each animal’s homeostatic setpoint; sleep-dominated windows (awake *<* 90%) were excluded. For the trajectory display (Fig. 1F), each animal’s wake *d*_2_ samples were plotted against time since the start of baseline and smoothed with a 6 h Gaussian kernel (±3 h); although the abscissa advances in clock time, only wake samples contributed and sleep periods contain no data points.

To test whether the setpoint is a durable individual trait, we partitioned the variance of the 5 min *d*_2_ estimates. The single-window reliability was the one-way random-effects intraclass correlation ICC(1, 1) (animal identity as the grouping factor); the reliability of the per-animal mean was ICC(1*, k*), corrected for the autocorrelation of successive windows by computing the effective number of independent windows (*n/τ*, *τ* ≈ 5 windows). Because consecutive windows are autocorrelated, the between-animal effect was tested by a one-way ANOVA on independent 3 h blocks with a label-permutation null, and reproducibility was corroborated by a split-half test–retest correlation (setpoint from the first vs. second non-overlapping half of baseline) with the Spearman–Brown full-length correction. Operational separability of animals was assessed by (i) the pairwise area under the ROC curve (AUC) for discriminating each pair of animals from single-window *d*_2_, related to the difference in setpoints by Spearman correlation, and (ii) an 18-way cross-validated linear-discriminant classifier of animal identity from *d*_2_, evaluated against a label-permutation null.

### Prey-capture scoring and learning-rate quantification (Fig. 1H, S5)

#### Capture-time scoring

The time to capture a red runner on each trial was scored by an expert rater. A hunt began when the roach was dropped into the arena and ended when the mouse bit off the roach’s head; when head-biting was not observed, the end was scored when the mouse settled into a stationary eating posture. Capture times were resolved to the nearest second and bounded above at 1,800 s (the 30 min timeout), with the exception of the pre-trial roach, which was available overnight; the pre-trial elapsed time is the true end − start interval and capture-vs-failure was read from whether the cockroach was dead or consumed at lights on. Capture times were log-transformed for all analyses.

#### Learning-rate quantification

Prey-capture learning speed was estimated from trial-by-trial capture times. Because learning can be conceptualized either as continuous improvement toward an asymptote^112,113^ (exponential model) or as skill acquisition followed by a plateau^114^ (breakpoint model), we extracted complementary metrics using two fits. For the **exponential model**, log capture time across trials *n* was fit to *y* = *Ae^−kn^* + *C* by nonlinear least squares, with the decay rate *k* constrained to [0, 0.5]; larger *k* indicates faster learning. To guard against timed-out trials driving the estimate, a second fit excluded timeout trials (≥ 1,800 s), yielding a second rate *k*_2_. For the **breakpoint model**, log capture time was fit to a piecewise-linear hinge with a flat tail, *y* = *α* − *β* min(*n* − *τ,* 0), in which performance improves with slope *β* up to breakpoint *τ* and is constant thereafter; *τ* was selected by exhaustive search over candidate trials (*τ* ≥ 3), minimizing the residual sum of squares. This yielded four duration-based metrics: breakpoint slope *β*, learned-trial number *τ*, exponential decay rate *k*, and exponential decay rate excluding failures *k*_2_. Because *τ* has the opposite sign convention, metrics were sign-corrected so that higher always indicates faster learning, standardized, and collapsed by principal-component analysis (PCA). The first principal component (PC1) explained 89.2% of the variance and served as the prey-capture learning-speed measure (Fig. S5F). Metric concordance was confirmed by Kendall’s *W*.

#### Early- and late-phase controls

Day-1 learning (Fig. 1G) used the median capture time over each animal’s first hunting day. Naive ability (Fig. S5C) used the first hunting trial, and pre- learning behavior (Fig. S5B) used the overnight pre-trial roach (continuous latency among animals that caught it; a binary caught/failed outcome otherwise). Asymptotic performance was indexed by the median capture time over each animal’s final ten trials, the fitted exponential asymptote *C*, and the mean late-phase latency; in-task locomotion (mean, peak, and total DLC velocity during hunts) was tested as an athleticism control.

### Ladder-cross preprocessing and behavioral quantification (Fig. 2)

#### Touch preprocessing (**Fig. 2B**)

Mice traversed the ladder in both directions; to make crosses directionally comparable, rungs were re-indexed so that rung 1 was always the first rung on the entry side and rung 10 the exit side. Rung contacts were detected by the MPR121 capacitive sensors, which register paw contacts but not tail contacts. To suppress within-contact sensor jitter, two touches on the same rung separated by less than 50 ms were merged into one, under the assumption that an animal cannot physically touch, lift, and re-touch a rung within 50 ms. Only crosses shorter than 60 s were retained. All quantitative ladder measures were derived from the MPR121 touch time series; video was used only to generate human labels for the cross-type classifier and contributed no measurement.

#### Trot detection by hidden Markov model (**Fig. 2C-E**)

To quantify the emergence of trot-like coordination, we fit a per-animal categorical hidden Markov model (HMM) to the sequence of rung- touch events^115^. The touch series for a cross was binned into 10 ms samples and represented as a binary pattern over a 5-rung window on each side of the ladder (2×5 = 10 bits), yielding 2^10^ possible observation symbols; only touches lasting ≥ 100 ms were included. Five hidden states captured the canonical trot cycle: the two diagonal stances (front-left/back-right and front-right/back-left), the two transient three-paw configurations bridging them, and an “other” state absorbing non-trot patterns (stalls, rearing, sniffing). The emission matrix was initialized by mapping each binary pattern to the state whose paw configuration it most closely matched; the transition matrix was initialized to favor the cyclic trot progression (1 → 2 → 3 → 4 → 1; transition 0.2, self-transition 0.7, 0.1 into “other”). Each per-animal model was then refined with hmmlearn on that animal’s last 300 crosses labeled “walk” or “trot” by the XGBoost classifier (below), matching each model to its own late-session gait statistics. After fitting, the hidden-state sequence for every cross was decoded, and a cross was classified as a trot if the product of posterior probabilities along the decoded trot- phase path exceeded 0.8^8^ ≈ 0.167 (80% confidence at each of the ∼ 8 phase transitions of a typical cross).

#### Behavioral classification of ladder crosses

To stratify the diverse behaviors mice display on the ladder, we trained an XGBoost classifier^105^ to assign each cross to one of six mutually exclusive categories: u-turn, stall-and-sniff, vertical exploration (rearing), walk, trot, and bound; the walk/trot/bound gait definitions follow Bellardita and Kiehn^117^. The classifier was trained on 2,328 manually labeled crosses from 8 animals. For each cross, 33 scalar and binary features were extracted, including cross duration, number of rungs touched, touch-time covariance, and left– right limb symmetry. Labeled data were split 70*/*30 into training and held-out test sets; random over-sampling corrected class imbalance in the training set, and a multi-class classifier was fit (max depth 5, 200 estimators), achieving 85% balanced accuracy on held-out data. The manifold analyses (Fig. 2I-J; Fig. S8C-D) used these XGBoost gait labels.

#### Manifold embedding of ladder crosses (**Fig. 2I-J**)

To analyze ladder crosses at the population scale, we embedded them in a low-dimensional manifold. Because crosses vary in their number of touches and a rung may be touched multiple times (especially early in learning), fixed-length-vector methods (PCA, UMAP) are poorly suited. We instead treated each cross as a sequence of touch events, computed pairwise distances by sequence alignment, and embedded the distance matrix in two dimensions by multidimensional scaling (MDS)^55,118^. Each cross was converted to an ordered sequence of touches, each touch described by four attributes (start time relative to cross onset, duration, body side, rung position 1–10); only touches ≥ 100 ms and crosses with more than four such touches were used (touches per cross: mean ± s.d. = 10.61 ± 5.14; range 5–150). Pairwise distances used Needleman–Wunsch (NW) global alignment^119^: the substitution cost between two touches was the sum of a 2-point penalty for opposite sides, the absolute rung-position difference, twice the relative duration difference, and four times the relative start-time difference; insertions/deletions incurred a gap cost of 16. MDS was run on the resulting per-animal distance matrix with 10 random initializations. For Fig. 2J and S8C-D, each animal was restricted to its first 200 crosses, and we computed the convex-hull area of the full embedding (the *all* -manifold area) and of the walk/trot subset (the *walk/trot* -manifold area); their ratio is the exploration/consolidation index (Fig. 2J).

Because metric MDS reproduces the absolute values of the input distances rather than their ranks, and all pairwise distances were computed with the same NW cost function and parameters, the resulting convex-hull areas share a common scale (*NWdistance*^2^) across animals. This makes cross-animal comparison of manifold size valid without post-hoc normalization. Scale-normalizing methods such as UMAP would not support this comparison, as they standardize local neighborhood structure and produce embeddings on an arbitrary per-animal scale.

#### Ferret plasticity and ***d*_2_** (Fig. 3)

For each ferret and exposure block, baseline *d*_2_ was computed from the assessment epoch immediately preceding exposure (above), and plasticity was the change in cell-averaged 1 − DCV across that block. This yielded up to three (*d*_2_, plasticity) points per animal. Points entered the correlation if the block passed the population-rate and spontaneous/evoked gates (non-control animals) or the rate gate alone (control animals), giving *N* = 33 blocks from 14 kits. Because the control-condition animals followed the same *d*_2_ →plasticity relation, including them tightened the fit; controls-included is the reported set. Because plasticity is an unbounded, signed change (unlike the floored, saturating mouse learning outcomes), the primary statistic here is Pearson *r* (with Spearman *ρ* and a permutation null as robustness checks). Pseudoreplication from multiple blocks per animal was addressed by a linear mixed-effects model, plasticity ∼ *d*_2_ + (1 | animal), and by collapsing to one point per animal (*N* = 14); the spike-time shuffle control (above) served as a negative control. The training-time increase in selectivity (Fig. 3B-C) was quantified by a linear mixed-effects model, direction selectivity ∼ exposure time + (1 | animal), over the motion animals.

#### Minimal recurrent model relating ***d*_2_** to learning rate (Fig. 3F-H)

Temporal credit assignment in a recurrent network trained by backpropagation through time is governed by a product of recurrent Jacobians that, for dynamics dominated by a single stable real mode *λ_∗_* ≡ *µ*, scales as *µ^k^* over a delay of *k* = *t* − *s* steps^67,68^. To obtain a closed-form relationship between distance to criticality and learning rate, we reduced the network to this dominant mode and analyzed learning of a delayed input–output association exactly.

#### Dominant-mode reduction and *d*_2_

Retaining only the slowest mode, the hidden dynamics reduce to the scalar recursion

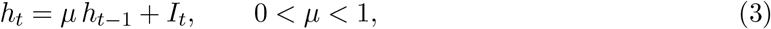

a first-order autoregressive (AR(1)) process. For an AR(*p*) model, 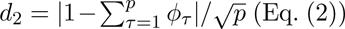 (Eq. (2)); for AR(1) (*p* = 1, *ϕ*_1_ = *µ*) this is simply

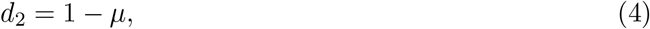

so proximity to criticality (*µ* → 1) corresponds exactly to *d*_2_ → 0. The credit-assignment gain over a delay of *k* steps is therefore

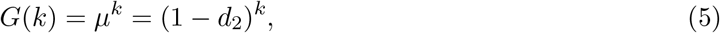

which decays geometrically with delay at a rate fixed entirely by *d*_2_ (Fig. 3F).

#### Exact learning dynamics

Each trial delivers a fixed input *I* at trial onset; the hidden state then decays freely, *h_τ_* = *µ^τ^ I*, and is read out *k* steps later, *h_k_* = *µ^k^I*, where *k* is the credit-assignment horizon of the task. A linear readout produces the output *o_m_* = *w_m_ h_k_* = *w_m_ µ^k^I* on trial *m*, with squared-error loss 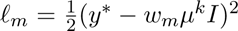 against a fixed target *y^∗^*. Gradient descent on *w* with step size *α* gives

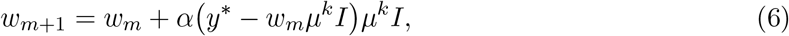

with unique optimum *w^∗^* = *y^∗^/*(*µ^k^I*) and loss floor *ℓ^∗^* = 0 for every network, independent of *d*_2_. Writing the error as *e_m_* = *w_m_* − *w^∗^*, the update contracts linearly,

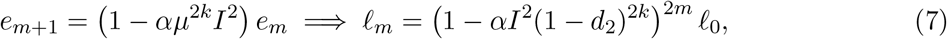

*\*where *ℓ*_0_ is the initial loss and the exponent 2*m* follows from the loss being quadratic in *e_m_*. Equation (7) gives the family of learning curves in Fig. 3G.

#### Trials to criterion

Defining learning as reaching a fixed loss tolerance *ℓ_M_*≤ *ɛ* and solving Eq. (7) for *M* gives the number of trials required,

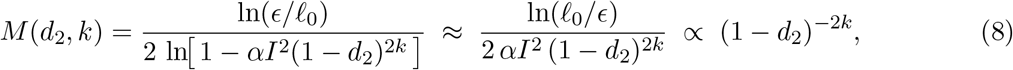

the small-step approximation (ln(1 − *z*) ≈ −*z*) holding whenever 0 *< z* ≡ *αI*^2^(1 − *d*_2_)^2*k*^ ≪ 1. Equation (8) is the surface in Fig. 3H; it decreases monotonically as *d*_2_ → 0 and steepens with the credit horizon *k*, so the penalty for being off-critical compounds with task difficulty.

#### Figure generation

All panels used *α* = 0.05, *I* = 1, *ℓ*_0_ = 1, and *ɛ* = 0.05. Fig. 3F plots the credit gain (Eq. (5)) against delay *k* ∈ [0, 60]; Fig. 3G plots the learning curves (Eq. (7)) against trial *m* ∈ [0, 500] at fixed *k* = 10; both show eight values of *d*_2_ evenly spaced over [0.05, 0.25]. Fig. 3H plots log_10_ *M* (Eq. (8)) over *d*_2_ ∈ [0.001, 0.30] and *k* ∈ [10, 100]. To avoid catastrophic cancellation when *z* is small, ln(1 − *z*) was evaluated as log1p(−*z*).

#### Human ***d*_2_** versus IQ (Fig. 4)

The analysis was between-patient: one *d*_2_ per patient versus one full-scale IQ, paralleling the mouse learning-rate design. A window-level mixed model with a patient random intercept is degenerate here because IQ is constant within patient (the random intercept absorbs the entire between-patient effect), so it was not used. The primary model was a partial correlation of *d*_2_ with IQ adjusting for demographic covariates (sex, age, race/ethnicity, and handedness), which are uncorrelated with *d*_2_ and therefore act as suppressors; the zero-order Pearson correlation is also reported. Significance of the adjusted effect was assessed by a Freedman–Lane residual-permutation test. Model form was checked by BIC over monotonic candidates (linear, exponential, power, reciprocal), which selected linear. Robustness was established by a 16-way aggregator sweep (window-to-patient collapse), leave-one-patient-out, and a clinical-sensitivity model additionally adjusting for autism, seizure burden, and resection status (which are entangled with *d*_2_ and attenuate the estimate; reported as a supplement, not the primary).

#### Baseline control-feature (specificity) panels (Fig. 1J; Figs. 2E,H; Fig. 3E; Fig. 4E)

To test whether *d*_2_’s predictive power is specific, in each figure *d*_2_ was competed against a panel of additional baseline features, each entered as a standalone univariate predictor of the same outcome. For mice, the panel comprised 20 control features spanning spike-rate/regularity statistics (mean RSU firing rate; CV, CV2, LV, LVR; Fano factor at 100 and 200 ms; Lempel–Ziv complexity of the 15 ms-binned population train), inter-spike-interval statistics (min, max, median, mode, and s.d. of the ISI), normalized LFP band power (delta, theta, alpha) and band ratios (gamma/alpha, gamma/beta), and animal-level covariates (DLC movement, wake proportion); with *d*_2_ this is 21 neural/behavioral features. For ferrets, only the 13 spike-train–computable features were available (behavioral and LFP features have no anesthetized equivalent). For humans, *d*_2_ was competed against 13 *amplitude-invariant* spectral features only—relative band powers, band ratios (*θ/β*, *γ/θ*, high-*γ/θ*), spectral entropy, and the envelope coefficient of variation—because absolute sEEG amplitude is set by non-physiological factors (electrode geometry, impedance, reference, gain) and is not comparable across patients; absolute-amplitude features (signal std/var/RMS, absolute and log band powers, envelope mean/std/var) form a single collinear nuisance factor and were excluded.

Each panel bar is the univariate effect size against the figure’s outcome, using the statistic native to that figure: Spearman *ρ*^2^ for the mouse learning outcomes (a bounded, saturating quantity), Pearson *r*^2^ for ferret plasticity (an unbounded signed change), and the demographic covariate-adjusted partial *r*^2^ for human IQ. Significance was read against a permutation null built by shuffling the outcome relative to each feature. Because the *ρ*^2^ null under the null hypothesis depends only on *n* (continuous metrics, no ties), a single large shared permutation null (200,000 draws) was built per *n* and used for both every feature’s permutation *p* and the two reference lines drawn on each panel: the null median (chance) and the 95th percentile (the per-test *p <* 0.05 boundary).

#### No multiple-comparisons correctio

No multiple-comparisons correction was applied to the specificity panels. These panels are confirmatory of a *negative* prediction—that the effect should be specific to *d*_2_—not discovery tests. A spurious hit in a control feature can only *weaken* the specificity claim, never strengthen it, so FDR/Bonferroni would perversely penalize thoroughness (adding arbitrary controls would lower *q* for the primary without changing its predictive power). Significance is therefore read against the chance and 95th-percentile reference lines and the permutation null.

#### Relationship of *d*_2_ to elementary signal statistics (Fig. 1D; Fig. S2)

To test whether *d*_2_ merely re-expresses a simpler feature such as variance or slow-frequency power, we characterized the full space of stationary second-order autoregressive [AR(2)] processes—the lowest-order model that supports the scale-invariant dynamics used to define criticality^40^. AR(2) processes are specified by (*ϕ*_1_*, ϕ*_2_*, σ*^2^), with the stationary region forming a bounded triangle in the (*ϕ*_1_*, ϕ*_2_) plane and the critical family lying along *ϕ*_1_ + *ϕ*_2_ = 1; there, 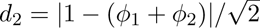 quantities of interest have closed forms over this plane: the stationary variance is

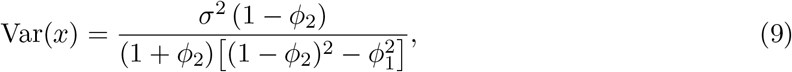

and the power spectral density is 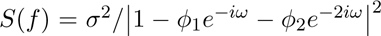 with *ω* = 2*πf* Δ*t*. To match the *d*_2_ analysis, spectral quantities used the same bin width, Δ*t* = 15 ms, with relative delta power defined as the fraction of total power in 0.5–4 Hz. Evaluated over the entire stable region, *d*_2_ increases monotonically with distance from the critical line, whereas variance and slow-frequency spectral content do not, and each conventional descriptor’s relationship to *d*_2_ is local—its sign and magnitude shift, and can invert, across the plane. Thus no single statistic tracks the critical boundary coherently across animals the way *d*_2_ does.

#### Single-neuron and population control metrics

The control-panel features were computed from the same RSU populations and 5 min wake windows used for *d*_2_. The Fano factor was computed per neuron and averaged across neurons to give one value per animal per window. Inter-spike-interval regularity metrics (CV, CV2, LV, LVR) were likewise restricted to RSUs, computed per neuron, and averaged per window^23^. Population Lempel–Ziv (LZ) complexity followed the binarization scheme of Abásolo et al.^120^: population spike trains were converted to binary sequences at 500 Hz (2 ms bins, 1 if ≥ 1 spike, else 0) at the 15 ms binning matched to *d*_2_, and LZ complexity was estimated from these binary population sequences. Normalized LFP band powers and band ratios were computed from the baseline LFP over the same windows.

### Statistics

Analyses were performed in Python (numpy, scipy, statsmodels, scikit-learn, hmmlearn, xgboost). Linear mixed-effects models (ferret plasticity and selectivity-vs-time) were fit with statsmodels (mixedlm, restricted maximum likelihood), with fixed-effect significance assessed by the Wald *z* test.

### Primary statistics by figure

For the mouse learning outcomes, the primary statistic is Spearman’s *ρ* (rank), because learning outcomes are bounded and saturating (capture time cannot fall below zero and decays to a floor); all mouse specificity panels square Spearman *ρ*^2^. For ferret plasticity, an unbounded signed change, the primary statistic is Pearson’s *r*, with Spearman *ρ* as a robustness check; a linear mixed-effects model with animal as a random effect provides the inferential *p*-value against block pseudoreplication. For the human analysis, the primary statistic is the demographic-adjusted partial correlation. Pearson/linear fits and BIC model-selection fits are reported alongside rank statistics where relevant but are not the headline result.

### Permutation nulls

For key correlations we report a permutation *p*-value obtained by shuffling one variable relative to the other, recomputing the correlation under each shuffle, and taking the proportion of shuffled correlations whose absolute value met or exceeded the observed value. Nulls used 5,000 shuffles for individual correlations and 200,000 for the shared specificity-panel nulls. The human adjusted effect used a Freedman–Lane residual permutation.

### Model selection

Where a functional form was fit to a scatter (e.g., the ladder exploration/ consolidation ratio, Fig. 2; the human *d*_2_–IQ relationship, Fig. 4), the model was selected by the Bayesian Information Criterion (BIC) over a set of *monotonic* candidates (linear, exponential, reciprocal, power); quadratic and other non-monotonic models were excluded, as a U-shape has no justification under the criticality hypothesis and overfits at these sample sizes. For the ladder ratio, an unbounded and magnitude-meaningful quantity, BIC was scored in log space (minimizing squared error in log_10_) to prevent a single high-leverage animal from dominating the raw-space fit, and Pearson *r* on log–log axes is reported; BIC selected a power law.

### Multiple comparisons

No multiple-comparisons correction was applied to the specificity panels, for the reasons given above (they are confirmatory of a negative prediction). Reported *p*-values are two-tailed and, unless noted, uncorrected.

### Data availability

Source data and derived tables supporting the findings of this study are available from the corresponding authors upon reasonable request. Human sEEG data are coded clinical data and are subject to the governing IRB and data-use agreements. Ferret data (Ritter et al.^60^) are available from S.D.V.H.

### Code availability

Analysis code is available at https://github.com/hengenlab.

---

**Figure S1:**
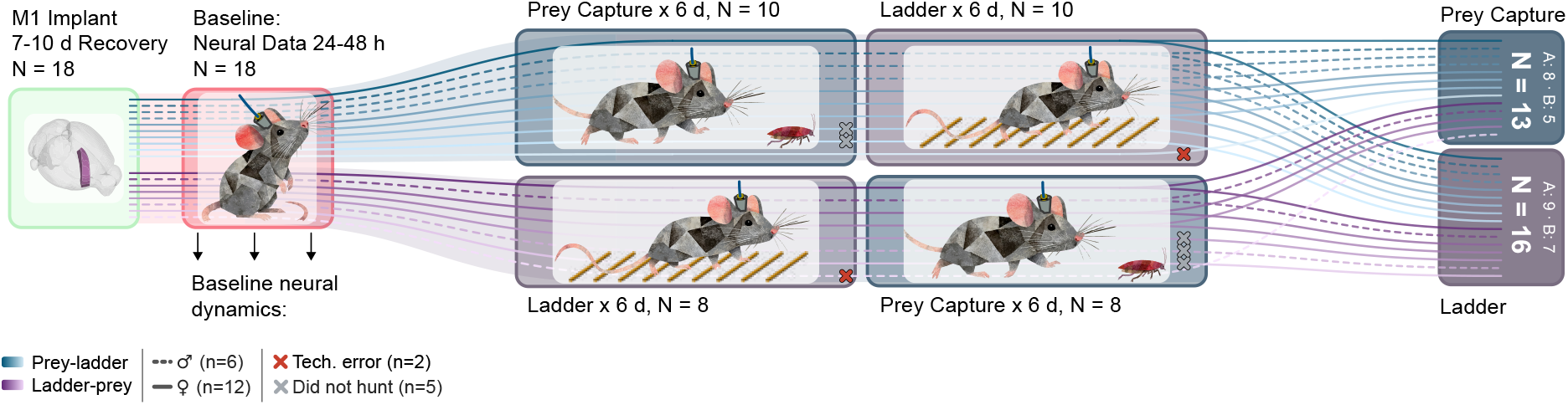
Experimental pipeline and per-animal task assignment. Eighteen mice (6 male, 12 female) received chronic right M1 microelectrode implants, recovered for 7–10 days, and underwent 24–48 h of task-free baseline neural recording before completing two counterbalanced 6-day behavioral tasks: prey capture (roach hunting, ~3 h/morning) and ladder crossing (continuously available in the homecage). Ten animals performed prey capture first (Group A, blue) and eight performed ladder crossing first (Group B, purple); solid lines trace individual females and dashed lines trace individual males across the pipeline. Downstream analysis datasets (right) comprise 16 animals with valid ladder-crossing data and 13 confirmed roach hunters; 11 animals contribute to both. Exclusions are marked with ×: two animals lost ladder data to equipment failure but retained valid hunting data (red, “technical ladder failure”), and five animals never hunted a roach and were excluded from the prey-capture analysis only (gray, behavioral non-consolidation unrelated to *d*_2_).

**Figure S2:**
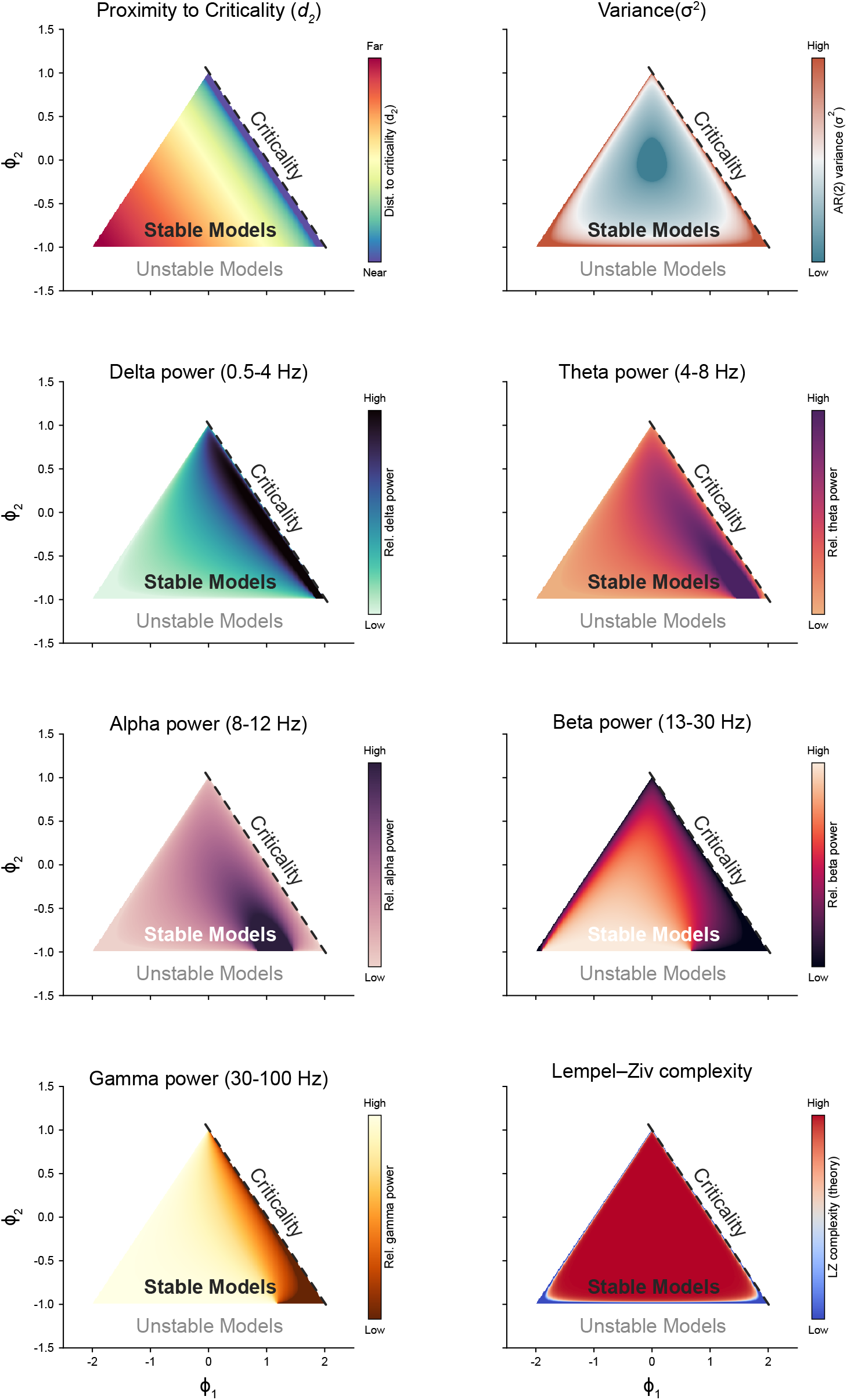
Features of neural activity exhibit distinct patterns when considered across a space of possible dynamics. Each panel maps a candidate metric across the stable region of the AR(2) parameter space (*ϕ*_1_–*ϕ*_2_ plane; critical boundary *ϕ*_1_ + *ϕ*_2_ = 1, dashed), colored by that metric’s value. Unlike *d*_2_ (top left), which increases monotonically with distance from the critical line everywhere in the space, other comparator descriptors—variance (*σ*^2^), relative band power (*δ*, *θ*, *α*, *β*, *γ*), and Lempel–Ziv complexity—tracks *d*_2_ only locally, with a relationship whose sign and magnitude shift across the plane.

**Figure S3:**
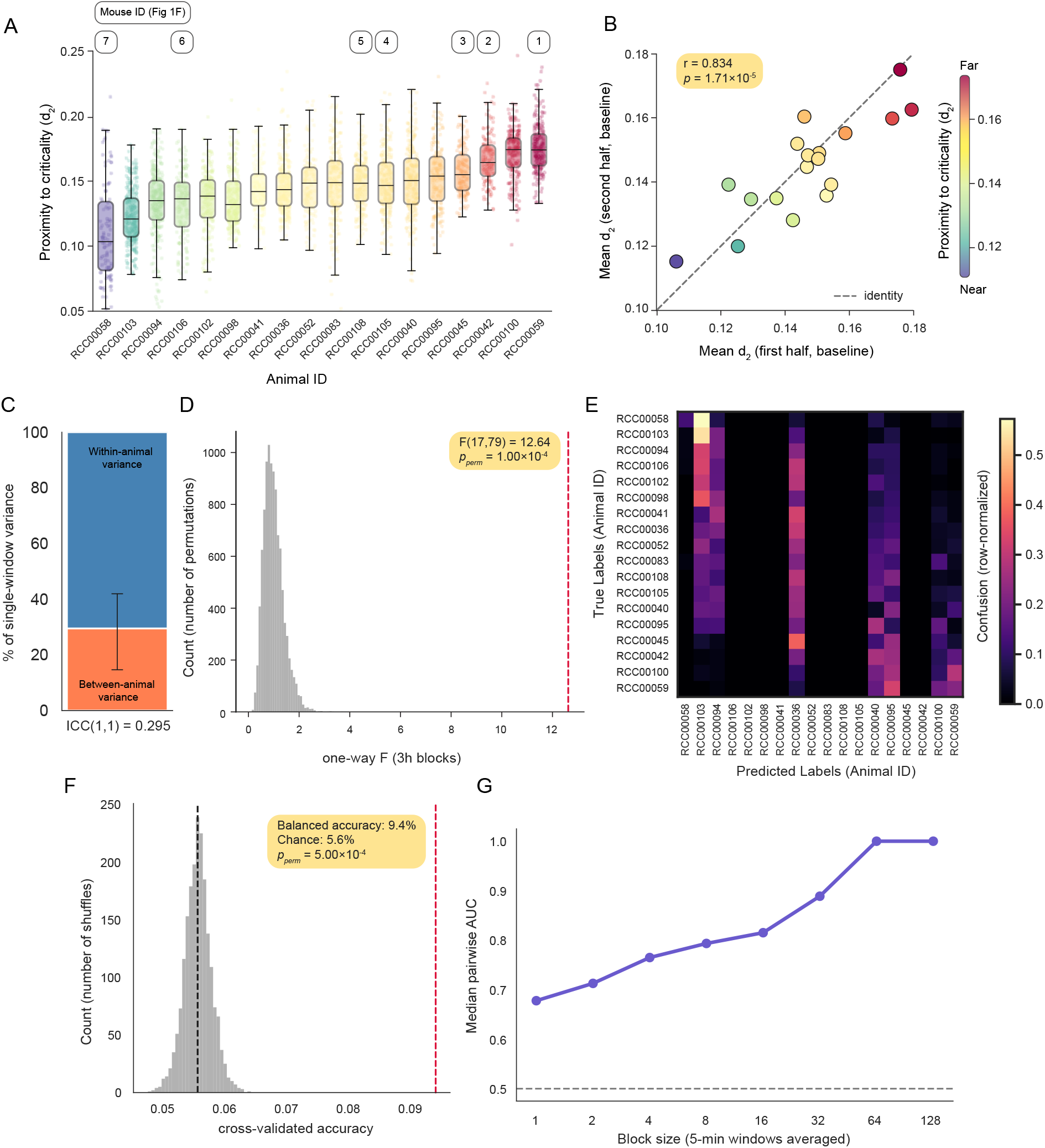
*d*_2_ is stable and discriminable as an individual setpoint. **(A)** Per-animal distribution of baseline *d*_2_ (300-s windows, M1, wake), animals ordered by mean *d*_2_; boxes show median and IQR, points are individual windows, color encodes mean *d*_2_ (*N* = 18). **(B)** Split-half test–retest: mean *d*_2_ from first vs. second half of each animal’s baseline (Pearson *r* = 0.834, *r*^2^ = 0.695, *p* = 1.71 × 10*^−^*^5^). **(C)** Variance decomposition of single-window *d*_2_: between-animal variance is only 29.5% (ICC(1,1)= 0.295, 95% CI [0.147, 0.420]). **(D)** One-way ANOVA for animal identity (F(17,79)= 12.64) falls far outside a 3-h-block label-shuffle permutation null (0/10,000 permutations observed, *p*_perm_ *<* 10*^−^*^4^), confirming the animal effect is not noise. **(E)** Confusion matrix from 18-way linear discriminant classification of animal identity from single- window *d*_2_ (2-fold interleaved cross-validation within animal; row-normalized). **(F)** Balanced classification accuracy against a label-shuffle null (2,000 shuffles). **(G)** Pairwise discriminability (cross-validated AUC) rises as 5-min *d*_2_ windows are averaged into longer blocks, from a median AUC of 0.68 at one window to 1.00 by ∼5.3 h of integration (64 windows).

**Figure S4:**
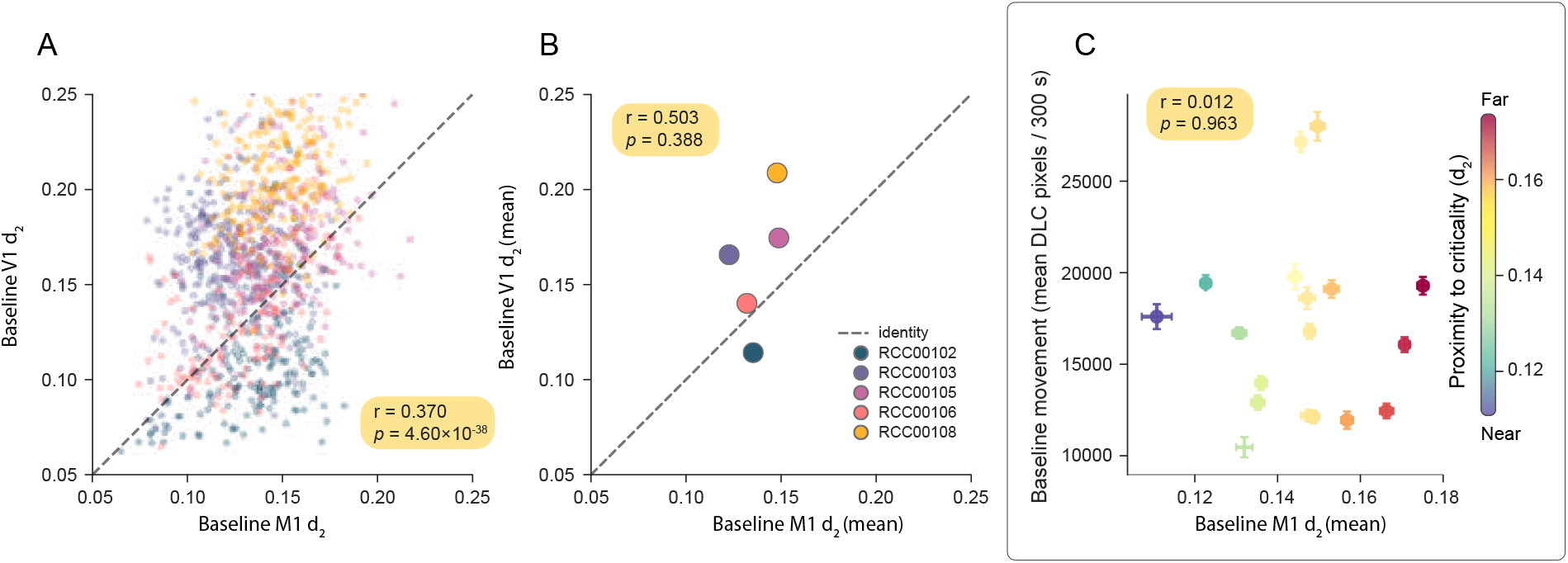
Baseline proximity to criticality is not a global, physiological set-point, nor is it a trivial proxy for baseline motor vigor. **(A)** For *N* = 5 mice where concurrent baseline data were recorded from M1 and V1, there is a weak correlation between M1 and V1 *d*_2_ when all non-overlapping epochs are considered (Pearson *r* = 0.37, *p* = 4.6 × 10*^−^*^38^, *n* = 1131 blocks of wake). **(B)** When the mean *d*_2_ set-points for M1 and V1 from these same mice are considered, there is no correlation (Pearson *r* = 0.50, *p* = 0.39, *N* = 5). **(C)** When baseline M1 *d*_2_ is considered as a predictor for the amount of running an animal does in the homecage, there is no correlation (Pearson *r* = 0.01, *p* = 0.96; Spearman *ρ* = − 0.05, *p* = 0.84; *N* = 18 animals).

**Figure S5:**
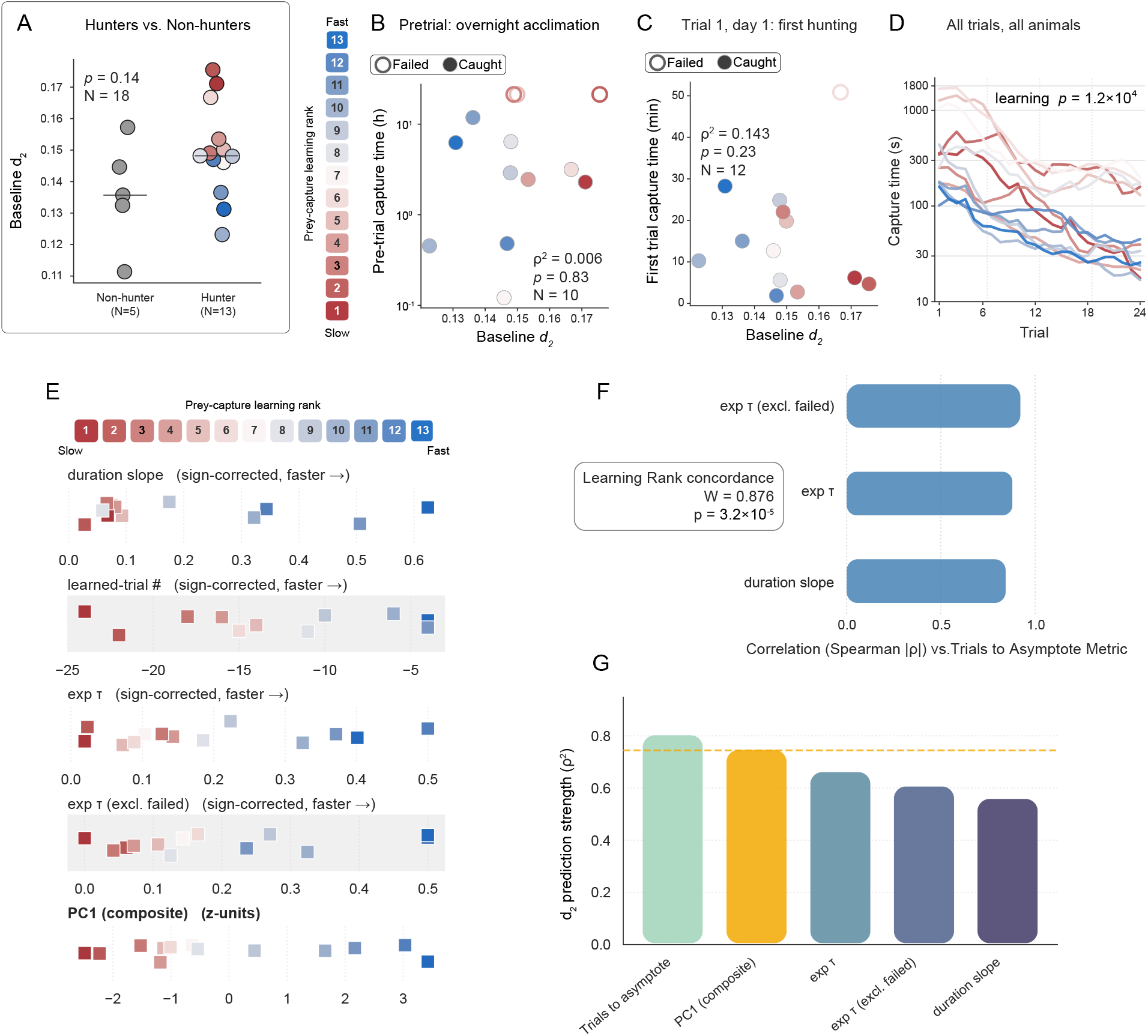
Animals show learning in a prey-capture paradigm, and initial performance is not predicted by proximity to criticality. **(A)** Baseline *d*_2_ does not differ between animals that hunted (Hunter, *N* = 13) and animals that never made a successful strike (Non-hunter, *N* = 5; Mann–Whitney *U* = 48.0, *p* = 0.14). Non-hunters are excluded from B–G as behaviorally non-interpretable (motivational or sensory, not slow learning), not because of their *d*_2_. **(B)** Capture time on the overnight acclimation roach, before any learning trial, is unrelated to baseline *d*_2_ (*N* = 10 that caught it; Spearman *ρ*^2^ = 0.006, *p_perm_* = 0.83); open circles never captured this roach. **(C)** Capture time on the first real trial (day 1, trial 1) is likewise unrelated to baseline *d*_2_ (*N* = 12 caught; *ρ*^2^ = 0.143, *p_perm_* = 0.23), so *d*_2_ does not track innate hunting ability at first exposure. **(D)** Every animal improves over the 24-trial course regardless of *d*_2_ (6-trial rolling-mean capture time; 13/13 animals show a negative within-animal Spearman correlation between capture time and trial number, sign test *p* = 1.2 10*^−^*^4^) – the *d*_2_ relationship emerges only once trials accumulate. **(E)** Distributions of the four constituent duration-learning metrics (duration slope, trials- to-asymptote, exponential time constant *τ*, and *τ* excluding failed hunts, each sign-corrected so higher = faster learner) and their first principal component (PC1, the composite learning score used throughout), animals colored by PC1 rank – the four metrics order animals concordantly. **(F)** Pairwise rank concordance (Spearman *ρ*) of each metric against the reference (trials-to-asymptote) is high (Kendall’s *W* = 0.876, *p* = 3.2 10*^−^*^5^), so the composite is not built from disagreeing metrics. **(G)** Baseline *d*_2_’s predictive power (Spearman *ρ*^2^) for each metric and for PC1 (gold); PC1 ranks 2nd of five for *d*_2_ prediction (*ρ*^2^ = 0.744), mid-range among its own constituents rather than an artificially inflated combination.

**Figure S6:**
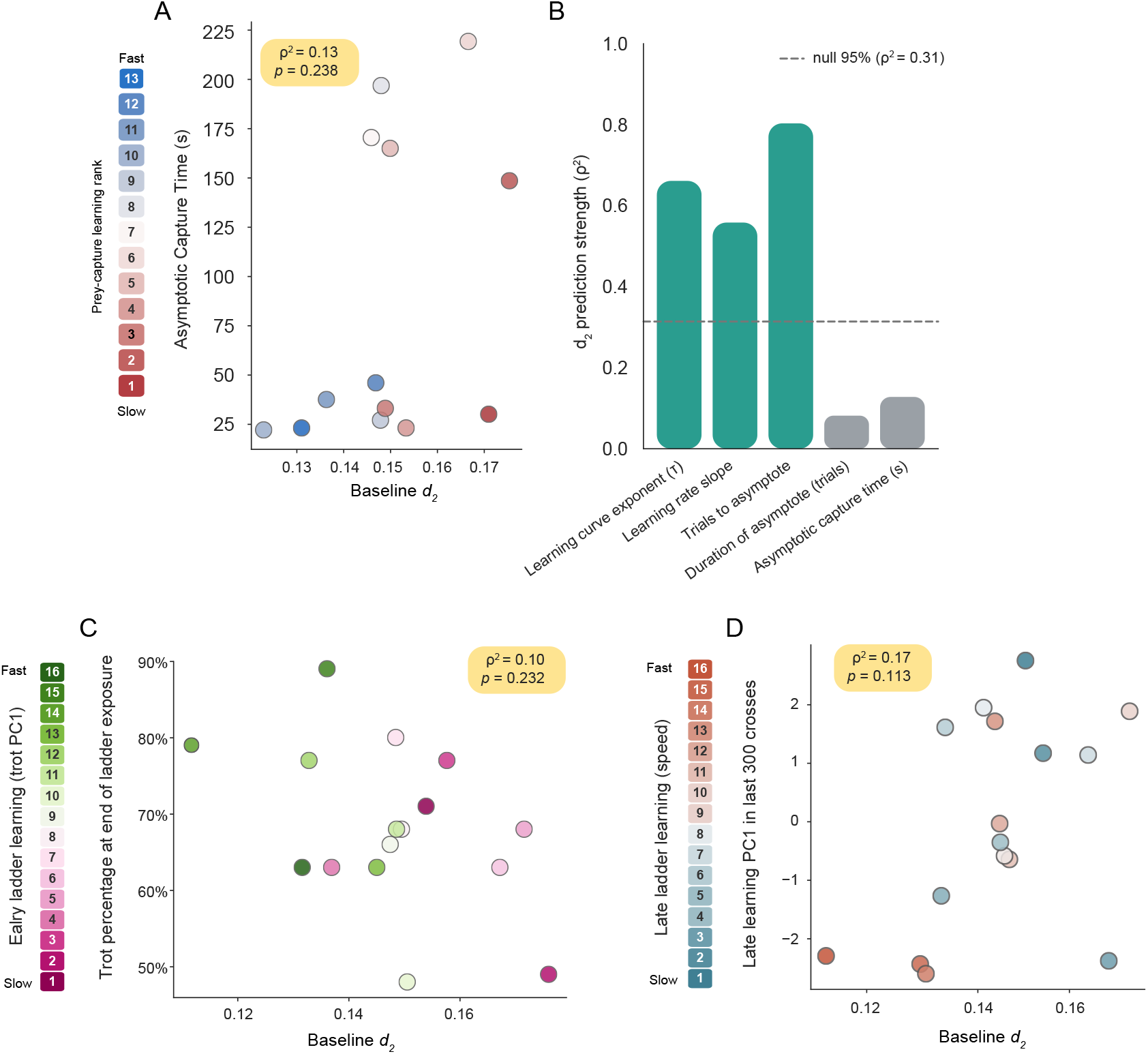
Proximity to criticality does not predict behavioral asymptote in prey-capture or ladder learning. **(A)** There was no correlation between baseline *d*_2_ and median capture time across each animal’s final 10 trials (its behavioral asymptote/ceiling) (Spearman *ρ* = 0.36, *ρ*^2^ = 0.13, *p_perm_*= 0.234; *N* = 13 hunters). **(B)** Compared to its prediction strength for behavioral-asymptote measures, baseline *d*_2_ was far more predictive of learning-*rate* measures: three rate metrics (exponential decay constant *k*: *ρ*^2^ = 0.66, *p_perm_* = 0.001; duration slope: *ρ*^2^ = 0.56, *p_perm_* = 0.005; learned-trial number: *ρ*^2^ = 0.80, *p_perm_* = 0.0002) all exceeded two asymptote metrics (asymptote *C*: *ρ*^2^ = 0.08, *p_perm_*= 0.341; median last-10 capture time: *ρ*^2^ = 0.13, *p_perm_* = 0.234), *N* = 13 for all. The dashed *null* line marks the 95th percentile of a permutation null for Spearman *ρ*^2^ (animal identity shuffled 5,000 times, matched to *N* = 13) — i.e., the effect size a metric would need to exceed by chance alone to reach an uncorrected *p <* 0.05 (*ρ*^2^ = 0.31); asterisks mark the metrics clearing this threshold. **(C)** Baseline *d*_2_ did not predict the endstage (ceiling) proportion of trots among the first 100 crossings of each animal’s final 300 ladder crossings (Spearman *ρ* = − 0.32, *ρ*^2^ = 0.10, *p* = 0.232; Pearson *r* = − 0.46, *r*^2^ = 0.21, *p* = 0.074; *N* = 16). **(D)** Baseline *d*_2_ likewise did not predict an endstage speed-milestone composite (PC1 of percentile on-ladder-duration milestones over the final 300 crossings; 99.6% variance explained) (Spearman *ρ* = +0.41, *ρ*^2^ = 0.17, *p* = 0.113; Pearson *r* = +0.45, *r*^2^ = 0.20, *p* = 0.083; *N* = 16).

**Figure S7:**
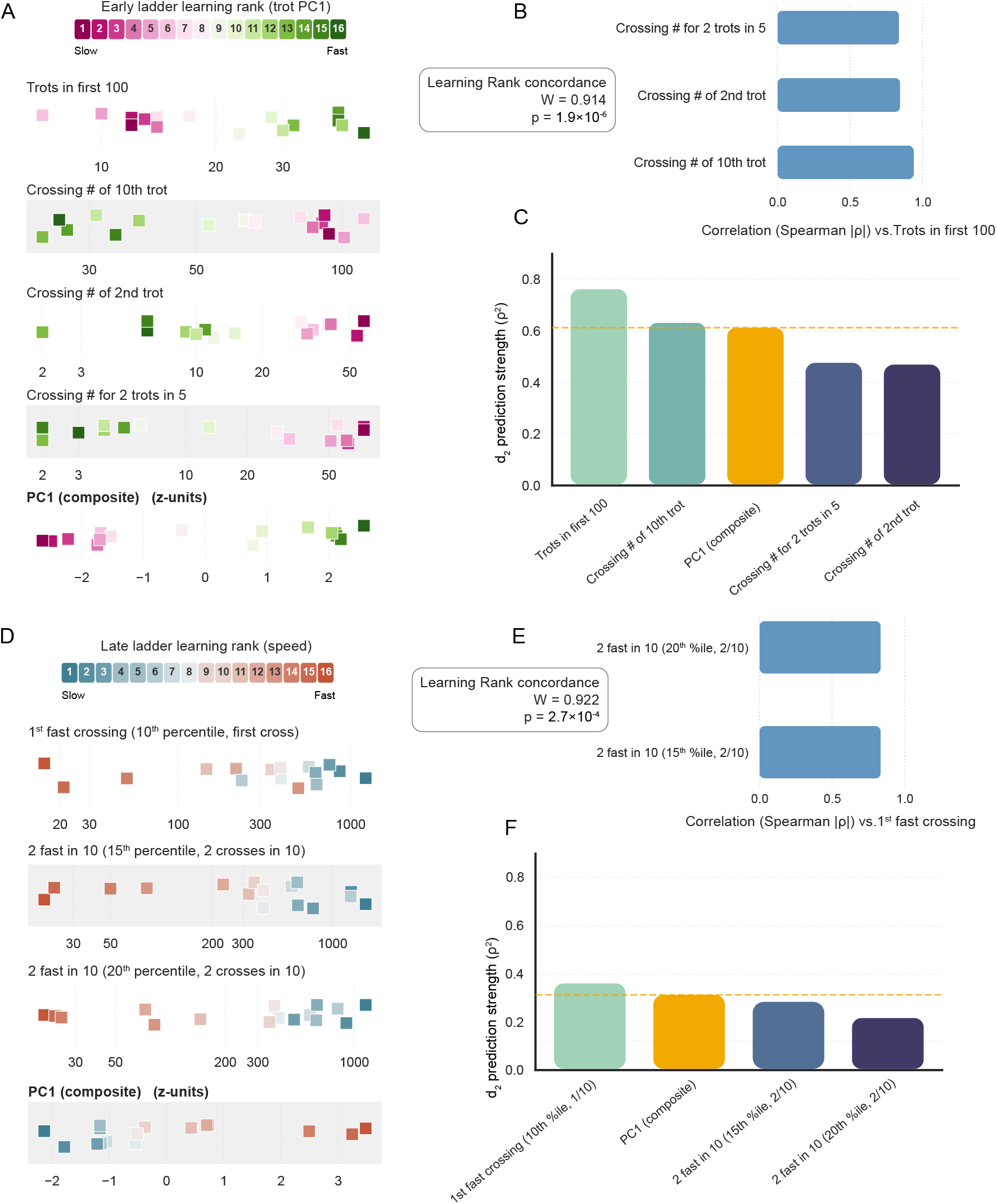
Animals learn and refine a repetitive motor sequence as they learn to cross a ladder. (A–C) Animals (*N* = 16) vary in how fast they learn to trot on the ladder. **(A)** Distributions of the four constituent trot-learning metrics (trots in the first 100 interactions, crosses to 10^th^ trot, crosses to 2^nd^ trot, and crosses until 2 trots occur within a 5-cross sliding window) and their first principal component (PC1, the composite learning score used throughout); animals colored by PC1 rank – the metrics order animals concordantly. **(B)** Pairwise rank concordance (Spearman *ρ*) of each metric against the reference (trots in the first 100 interactions) is high (Kendall’s *W* = 0.914, *p* = 1.9 10*^−^*^6^), so the composite is not built from disagreeing metrics. **(C)** Baseline *d*_2_’s predictive power (Spearman *ρ*^2^) for each metric and for PC1 (gold); PC1 ranks 3^rd^ of five (*ρ*^2^ = 0.61), mid-range among its own constituents rather than an artificially inflated combination. **(D–F)** Animals vary in how fast they approach an individual cross duration asymptote. **(D)** Distributions of the three constituent cross-duration-learning metrics (1^st^ cross reaching that mouse’s 10^th^ percentile for cross duration, crosses until 2 of 10 reach the 15^th^ percentile for speed, and crosses until 2 of 10 reach the 20^th^ percentile) and their first principal component (PC1); animals colored by PC1 rank – the metrics order animals concordantly. **(E)** Pairwise rank concordance (Spearman *ρ*) of each metric against the reference (crosses until the mouse reaches its 10^th^ percentile for speed) is high (Kendall’s *W* = 0.922, *p* = 2.7 10*^−^*^4^), so the composite is not built from disagreeing metrics. **(F)** Baseline *d*_2_’s predictive power (Spearman *ρ*^2^) for each metric and for PC1 (gold); PC1 ranks 2^nd^ of four (*ρ*^2^ = 0.31), mid-range among its own constituents rather than an artificially inflated combination.

**Figure S8:**
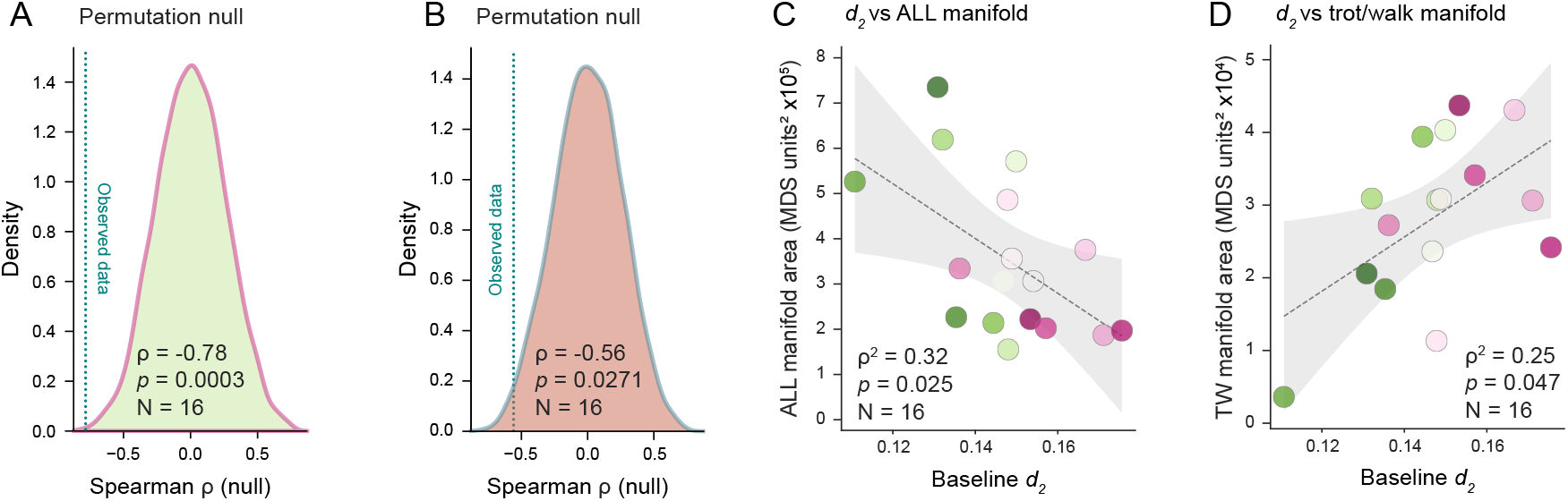
Mice learning the ladder show a robust prediction of early learning, late learning, and the geometric structure of behavior during learning. **(A)** Permutation null establishes that baseline *d*_2_ predicts early ladder learning (early-learning composite, PC1) far beyond chance: the observed Spearman correlation (*ρ* = −0.78, teal dashed line) falls outside the extreme tail of a null distribution built from 200,000 label-shuffled draws (*p*_perm_ = 0.0003, *N* = 16 animals). **(B)** The same permutation test applied to late ladder learning (speed-milestone composite, PC1) shows baseline *d*_2_ remains a significant predictor, albeit with a smaller effect size (*ρ* = −0.56, *p*_perm_ = 0.027, *N* = 16). **(C)** Baseline *d*_2_ also predicts the geometric structure of behavior during learning: animals closer to criticality (lower *d*_2_) traverse a larger overall movement repertoire, quantified as the area of the MDS-embedded behavioral manifold spanning all locomotor sub-types over the first 200 ladder crossings (*ρ*^2^ = 0.32, *p_perm_* = 0.025, *N* = 16). **(D)** Conversely, the manifold restricted to purposeful trot/walk locomotion is smaller in more-critical animals and grows as *d*_2_ increases (*ρ*^2^ = 0.25, *p_perm_* = 0.047, *N* = 16) — so animals near criticality explore a broad overall repertoire built around a comparatively tight trot/walk core. In **C–D**, dots are colored by each animal’s early-learning rank.

**Figure S9:**
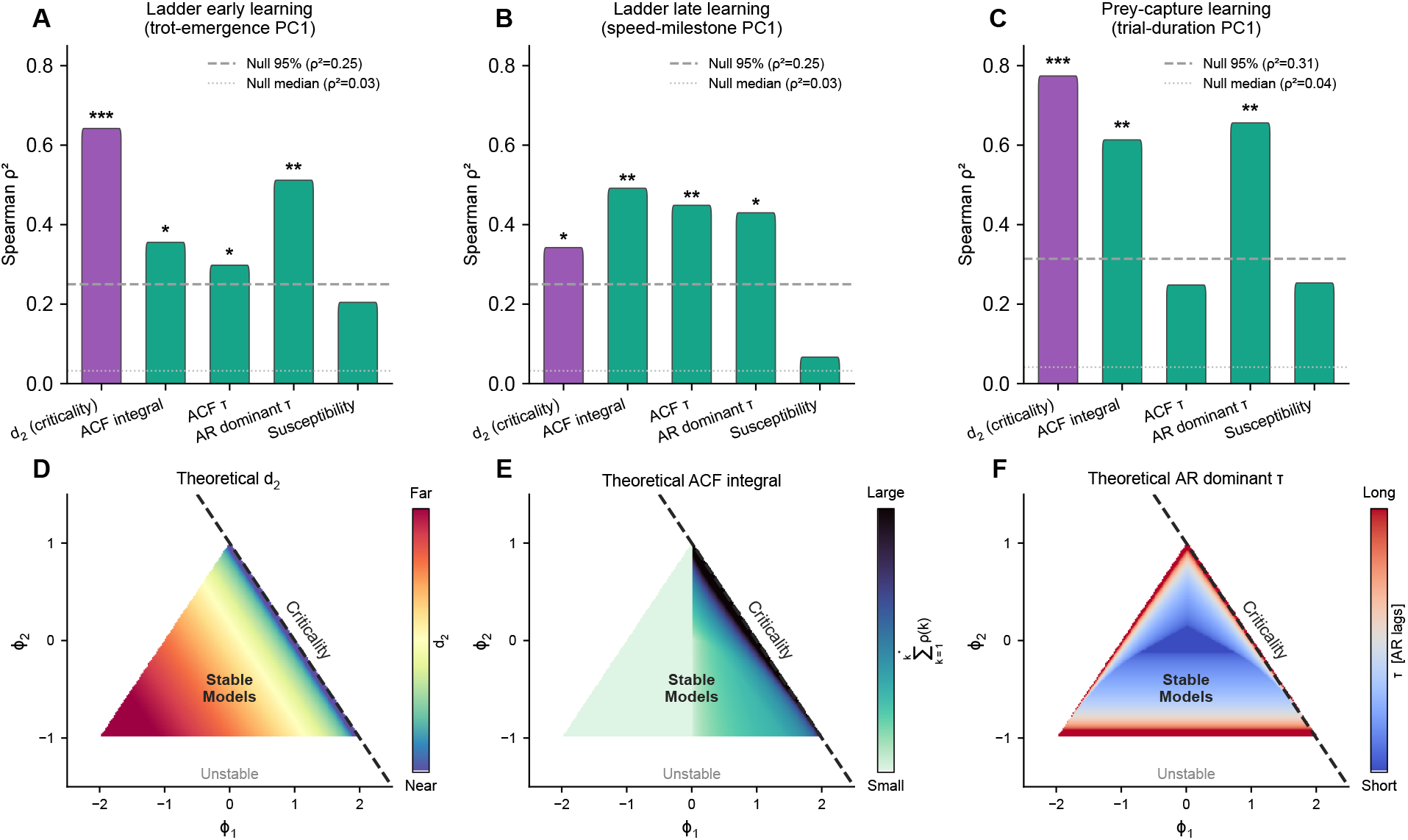
Autocorrelation timescales predict future learning and share dynamical origins with. *d*_2_ **in AR(2) space.** All metrics were derived from the 300 s wake baseline RSUs activity, using 15 ms population spike-count bins. *d*_2_ (purple) is identical to that used throughout the paper. **ACF integral** is the area under the population activity autocorrelation function (ACF) up to its first zero-crossing. The ACF was fit with *Ae^−t/τ^* + *B*, and the asymptotic floor *B* was subtracted. **ACF** *τ* is the exponential time constant from the same fit. **AR dominant** *τ* is derived from the same AR(10) model used to compute *d*_2_. AR dominant *τ* = − Δ*t/* ln *λ*_max_, where *λ*_max_ is the dominant eigenvalue of the AR(10) companion matrix (from Yule–Walker); this is the memory timescale implied by the fitted model. **Susceptibility** is *N* Var(*ρ_t_*), where *ρ_t_* is the fraction of recorded cells that fired at least once (binarized) in bin *t*, following Eq. 2 of^121^. **(A–C)** Both *d*_2_ and the autocorrelation measures (ACF integral, ACF *τ*, and AR dominant *τ*) predict ladder- and prey-capture learning, whereas susceptibility does not reach significance. **(D–F)** Theoretical behavior of each metric across the full space of stable AR(2) models (*ϕ*_1_–*ϕ*_2_ plane; dashed line, critical boundary *ϕ*_1_ + *ϕ*_2_ = 1). **(D)** The *d*_2_ triangle is identical to that in Fig. 1D. The susceptibility triangle is not shown, as it matches the “Variance *σ*^2^” heatmap in Fig. S2. **(E)** The ACF integral (sum of *ρ*(*k*) to the first zero-crossing) diverges at the critical line and falls to zero far from it. **(F)** The dominant autocorrelation timescale *τ* shows the same divergence at criticality.

**Supplementary Videos** Path: 2026_chopra_zhong/database/supplemental_videos

- Supplementary Video 1 (Early Crossing): ladder_early_crossing.mp4
- Supplementary Video 2 (trot, angle 1): ladder_trot_angel1.mp4
- Supplementary Video 3 (trot, angle 2): ladder_trot_angel2.mp4
- Supplementary Video 4 (trot at 0.25 x speed, angle 2): ladder_trot_0p25x_angle2.mp4
- Supplementary Video 5 (u-turn): ladder_uturn.mp4
- Supplementary Video 6 (vertical exploration): ladder_vertical_exploration.mp4

## Notes

### Competing Interest Statement

The authors have declared no competing interest.

### Summary of Updates

Fixed typos, corrected author order. We have also updated a statistic in Supplemental figure 5 where the incorrect p value was included for a permutation test (does not change statistical interpretation of the finding, but p value that was listed was incorrect).

### Funding

- National Institutes of Health: R01NS118442-01

## References

1. Darshil Doshi, Tianyu He, and Andrey Gromov. “Critical Initialization of Wide and Deep Neural Networks through Partial Jacobians: General Theory and Applications.” In: (Nov. 2021). arXiv: 2111.12143. url: http://arxiv.org/abs/2111.12143.

[2] Weixuan Liu, Xinyue Zhang, and Yuhan Helena Liu. “The Influence of Initial Connectivity on Biologically Plausible Learning.” In: arXiv (2024). doi: 10.48550/arxiv.2410.11164. eprint: 2410.11164.

[3] Rainer Engelken. “Gradient Flossing: Improving Gradient Descent through Dynamic Control of Jacobians.” In: arXiv (2023). doi: 10.48550/arxiv.2312.17306. eprint: 2312.17306.

[4] Samuel S Schoenholz et al. “Deep information propagation.” In: Conference paper of ICLR 2017 (2017), pp. 1–18. arXiv: arXiv:1611.01232v2.

[5] Arsham Ghavasieh, et al. “Toward a Physics of Deep Learning and Brains.” In: arXiv (2025). doi: 10.48550/arxiv.2509.22649. eprint: 2509.22649.

[6] Robert Legenstein and Wolfgang Maass. “Edge of chaos and prediction of computational performance for neural circuit models.” In: Neural Networks 20.3 (2007), pp. 323–334. issn: 0893-6080. doi: 10.1016/j.neunet.2007.04.017.

[7] Nils Bertschinger and Thomas Natschläger. “Real-Time Computation at the Edge of Chaos in Recurrent Neural Networks.” In: Neural Computation 16.7 (July 2004), pp. 1413–1436. issn: 1530-888X. doi: 10.1162/089976604323057443. url: http://dx.doi.org/10.1162/089976604323057443.

[8] David Dahmen et al. “Second type of criticality in the brain uncovers rich multiple-neuron dynamics.” In: Proceedings of the National Academy of Sciences of the United States of America 116.26 (2019), pp. 13051–13060. issn: 10916490. doi: 10.1073/pnas.1818972116.

[9] Guillermo B. Morales, Serena di Santo, and Miguel A. Muñoz. “Quasiuniversal scaling in mouse-brain neuronal activity stems from edge-of-instability critical dynamics.” In: Proceedings of the National Academy of Sciences of the United States of America 120.9 (2023), pp. 1–12. issn: 10916490. doi: 10.1073/pnas.2208998120. arXiv: 2111.12067.

[10] Keith B. Hengen and Woodrow L. Shew. “Is criticality a unified setpoint of brain function?” In: Neuron (June 2025). issn: 0896-6273. doi: 10.1016/j.neuron.2025.05.020. url: http://dx.doi.org/10.1016/j.neuron.2025.05.020.

[11] John M. Beggs. The Cortex and the Critical Point. Cambridge, MA: MIT Press, 2022. isbn: 9780262544030.

[12] John M. Beggs and Dietmar Plenz. “Neuronal Avalanches in Neocortical Circuits.” In: The Journal of Neuroscience 23.35 (Dec. 2003), pp. 11167–11177. issn: 1529-2401. doi: 10.1523/jneurosci.23-35-11167.2003. url: http://dx.doi.org/10.1523/JNEUROSCI.23-35-11167.2003.

[13] Benjamin Cramer et al. “Control of criticality and computation in spiking neuromorphic networks with plasticity.” In: Nature Communications 11.1 (2020). issn: 2041-1723. doi: 10.1038/s41467-020-16548-3. url: http://dx.doi.org/10.1038/s41467-020-16548-3.

[14] Simon Vock and Christian Meisel. Critical dynamics governs deep learning. 2025. doi: doi.org/ 10.48550/arXiv.2507.08527. arXiv: 2507.08527 [q-bio.NC].

[15] Clayton Haldeman and John M. Beggs. “Critical Branching Captures Activity in Living Neural Networks and Maximizes the Number of Metastable States.” In: Physical Review Letters 94.5 (2005), p. 058101. issn: 0031-9007. doi: 10.1103/physrevlett.94.058101.

[16] Woodrow L. Shew et al. “Information Capacity and Transmission Are Maximized in Balanced Cortical Networks with Neuronal Avalanches.” In: The Journal of Neuroscience 31.1 (Jan. 2011), pp. 55–63. issn: 1529-2401. doi: 10.1523/jneurosci.4637-10.2011. url: http://dx.doi.org/10.1523/JNEUROSCI.4637-10.2011.

[17] Erik D. Fagerholm et al. “Cortical Entropy, Mutual Information and Scale-Free Dynamics in Waking Mice.” In: Cerebral Cortex 26.10 (July 2016), pp. 3945–3952. issn: 1460-2199. doi: 10.1093/cercor/bhw200. url: http://dx.doi.org/10.1093/cercor/bhw200.

[18] Vidit Agrawal et al. “Robust entropy requires strong and balanced excitatory and inhibitory synapses.” In: Chaos: An Interdisciplinary Journal of Nonlinear Science 28.10 (2018), p. 103115. issn: 1054-1500. doi: 10.1063/1.5043429. eprint: 1804.05266.

[19] Kenneth G. Wilson. “Renormalization Group and Critical Phenomena. I. Renormalization Group and the Kadanoff Scaling Picture.” In: Physical Review B 4.9 (Nov. 1971), pp. 3174– 3183. issn: 0556-2805. doi: 10.1103/physrevb.4.3174. url: http://dx.doi.org/10.1103/PhysRevB.4.3174.

[20] Kenneth G. Wilson. “Renormalization Group and Critical Phenomena. II. Phase-Space Cell Analysis of Critical Behavior.” In: Physical Review B 4.9 (Nov. 1971), pp. 3184–3205. issn: 0556-2805. doi: 10.1103/physrevb.4.3184. url: http://dx.doi.org/10.1103/PhysRevB.4.3184.

[21] Kenneth G. Wilson. “Problems in Physics with many Scales of Length.” In: Scientific American 241.2 (1979), pp. 158–179. issn: 0036-8733. doi: 10.1038/scientificamerican0879-158.

[22] Zhengyu Ma et al. “Cortical Circuit Dynamics Are Homeostatically Tuned to Criticality In Vivo.” In: Neuron 104.4 (Nov. 2019), 655–664.e4. issn: 0896-6273. doi: 10.1016/j.neuron.2019.08.031. url: http://dx.doi.org/10.1016/j.neuron.2019.08.031.

[23] Y. Xu, et al. “Sleep restores an optimal computational regime in cortical networks.” In: Nature Neuroscience (2024), pp. 1–11.

[24] Christian Meisel et al. “Decline of long-range temporal correlations in the human brain during sustained wakefulness.” In: Scientific Reports 7.1 (Sept. 2017). issn: 2045-2322. doi: 10.1038/s41598-017-12140-w. url: http://dx.doi.org/10.1038/s41598-017-12140-w.

[25] Wesley P. Clawson et al. “Adaptation towards scale-free dynamics improves cortical stimulus discrimination at the cost of reduced detection.” In: PLOS Computational Biology 13.5 (May 2017). Ed. by Claus C. Hilgetag, e1005574. issn: 1553-7358. doi: 10.1371/journal.pcbi.1005574. url: http://dx.doi.org/10.1371/journal.pcbi.1005574.

[26] Woodrow L. Shew et al. “Adaptation to sensory input tunes visual cortex to criticality.” In: Nature Physics 11.8 (June 2015), pp. 659–663. issn: 1745-2481. doi: 10.1038/nphys3370. url: http://dx.doi.org/10.1038/nphys3370.

[27] Walter B. Cannon. The Wisdom of the Body. New York: W. W. Norton & Company, 1932.

[28] Enzo Tagliazucchi et al. “Criticality in Large-Scale Brain fMRI Dynamics Unveiled by a Novel Point Process Analysis.” In: Frontiers in Physiology 3 (2012). issn: 1664-042X. doi: 10.3389/fphys.2012.00015. url: http://dx.doi.org/10.3389/fphys.2012.00015.

[29] Sabrina A Jones et al. “Scale-free behavioral dynamics directly linked with scale-free cortical dynamics.” In: eLife 12 (Jan. 2023). issn: 2050-084X. doi: 10.7554/elife.79950. url: http://dx.doi.org/10.7554/eLife.79950.

[30] Antonio J. Fontenele et al. “Low-dimensional criticality embedded in high-dimensional awake brain dynamics.” In: Science Advances 10.17 (2024), eadj9303. doi: 10.1126/sciadv.adj9303.

[31] Leandro J. Fosque et al. “Two views of the brain are reconciled by a unifying principle of maximal information processing.” In: (Nov. 2025). doi: 10.1101/2025.11.25.690580. url: http://dx.doi.org/10.1101/2025.11.25.690580.

[32] Shervin Safavi et al. “Signatures of criticality in efficient coding networks.” In: Proceedings of the National Academy of Sciences 121.41 (Oct. 2024). issn: 1091-6490. doi: 10.1073/pnas.2302730121. url: http://dx.doi.org/10.1073/pnas.2302730121.

[33] Osame Kinouchi and Mauro Copelli. “Optimal dynamical range of excitable networks at criticality.” In: Nature Physics 2.5 (Apr. 2006), pp. 348–351. issn: 1745-2481. doi: 10.1038/nphys289. url: http://dx.doi.org/10.1038/nphys289.

[34] Paul Manuel Müller, et al. “Critical dynamics predicts cognitive performance and provides a common framework for heterogeneous mechanisms impacting cognition.” In: Proceedings of the National Academy of Sciences 122.14 (2025), e2417117122. issn: 0027-8424. doi: 10.1073/pnas.2417117122.

[35] Ziad Obermeyer, Jasmeet K Samra, and Sendhil Mullainathan. “Individual differences in normal body temperature: longitudinal big data analysis of patient records.” In: BMJ (Dec. 2017), j5468. issn: 1756-1833. doi: 10.1136/bmj.j5468. url: http://dx.doi.org/10.1136/bmj.j5468.

[36] Keith B. Hengen et al. “Firing Rate Homeostasis in Visual Cortex of Freely Behaving Rodents.” In: Neuron 80.2 (Oct. 2013), pp. 335–342. issn: 0896-6273. doi: 10.1016/j.neuron.2013.08.038. url: http://dx.doi.org/10.1016/j.neuron.2013.08.038.

[37] Keith B. Hengen et al. “Neuronal Firing Rate Homeostasis Is Inhibited by Sleep and Promoted by Wake.” In: Cell 165.1 (Mar. 2016), pp. 180–191. issn: 0092-8674. doi: 10.1016/j.cell.2016.01.046. url: http://dx.doi.org/10.1016/j.cell.2016.01.046.

[38] Cristopher M. Niell and Michael P. Stryker. “Highly Selective Receptive Fields in Mouse Visual Cortex.” In: The Journal of Neuroscience 28.30 (2008), pp. 7520–7536. issn: 1529-2401. doi: 10.1523/jneurosci.0623-08.2008. url: http://dx.doi.org/10.1523/JNEUROSCI.0623-08.2008.

[39] David F. Parks et al. “A nonoscillatory, millisecond-scale embedding of brain state provides insight into behavior.” In: Nature Neuroscience (July 2024). issn: 1546-1726. doi: 10.1038/s41593-024-01715-2. url: http://dx.doi.org/10.1038/s41593-024-01715-2.

[40] J. Samuel Sooter et al. “Defining and measuring proximity to criticality.” In: (Aug. 2025). doi: 10.1101/2025.08.03.668332. url: http://dx.doi.org/10.1101/2025.08.03.668332.

[41] Adele Diamond et al. “One size does not fit all: Assuming the same normal body temperature for everyone is not justified.” In: PLOS ONE 16.2 (Feb. 2021). Ed. by Henrik Oster, e0245257. issn: 1932-6203. doi: 10.1371/journal.pone.0245257. url: http ://dx.doi.org/10.1371/journal.pone.0245257.

[42] Melvin Khee-Shing Leow and Simon L Goede. “The homeostatic set point of the hypothalamuspituitary-thyroid axis – maximum curvature theory for personalized euthyroid targets.” In: Theoretical Biology and Medical Modelling 11.1 (Aug. 2014). issn: 1742-4682. doi: 10.1186/1742-4682-11-35. url: http://dx.doi.org/10.1186/1742-4682-11-35.

[43] Suzanne C. Segerstrom et al. “Variability and reliability of diurnal cortisol in younger and older adults: Implications for design decisions.” In: Psychoneuroendocrinology 49 (Nov. 2014), pp. 299–309. issn: 0306-4530. doi: 10.1016/j.psyneuen.2014.07.022. url:http://dx.doi.org/10.1016/j.psyneuen.2014.07.022.

[44] Rajiv Ranganathan, Simon Cone, and Brian Fox. “Predicting individual differences in motor learning: A critical review.” In: Neuroscience& Biobehavioral Reviews 141 (Oct. 2022), p. 104852. issn: 0149-7634. doi: 10.1016/j.neubiorev.2022.104852. url: http://dx.doi.org/10.1016/j.neubiorev.2022.104852.

[45] Peilun Song et al. “Multi-network Topology Underlying Individual Language Learning Success.” In: The Journal of Neuroscience (2026), e2205252026. issn: 1529-2401. doi: 10.1523/jneurosci.2205-25.2026. url: http://dx.doi.org/10.1523/JNEUROSCI.2205-25.2026.

[46] Wilson S. Geisler and Duane G. Albrecht. “Visual cortex neurons in monkeys and cats: Detection, discrimination, and identification.” In: Visual Neuroscience 14.5 (1997), pp. 897– 919. issn: 1469-8714. doi: 10.1017/s0952523800011627. url: http://dx.doi.org/10.1017/s0952523800011627.

[47] Shigeru Shinomoto, Keisetsu Shima, and Jun Tanji. “Differences in Spiking Patterns Among Cortical Neurons.” In: Neural Computation 15.12 (Dec. 2003), pp. 2823–2842. issn: 1530-888X. doi: 10.1162/089976603322518759.

[48] G. R. Holt et al. “Comparison of discharge variability in vitro and in vivo in cat visual cortex neurons.” In: Journal of Neurophysiology 75.5 (May 1996), pp. 1806–1814. issn: 1522-1598. doi: 10.1152/jn.1996.75.5.1806. url: http://dx.doi.org/10.1152/jn.1996.75.5.1806.

[49] Luca Cocchi et al. “Criticality in the brain: A synthesis of neurobiology, models and cognition.” In: Progress in Neurobiology 158 (Nov. 2017), pp. 132–152. issn: 0301-0082. doi: 10.1016/j.pneurobio.2017.07.002. url: http://dx.doi.org/10.1016/j.pneurobio.2017.07.002.

[50] Jordan O’Byrne and Karim Jerbi. “How critical is brain criticality?” In: Trends in Neurosciences 45.11 (Nov. 2022), pp. 820–837. issn: 0166-2236. doi: 10.1016/j.tins.2022.08.007. url: http://dx.doi.org/10.1016/j.tins.2022.08.007.

[51] Yahya Karimipanah, Zhengyu Ma, and Ralf Wessel. *New hallmarks of criticality in recurrent neural networks*. arXiv:1610.01217v2 [q-bio.NC]. 2016. doi: 10.48550/arXiv.1610.01217. arXiv: 1610.01217 [q-bio.NC].

[52] Risa Kawai et al. “Motor Cortex Is Required for Learning but Not for Executing a Motor Skill.” In: Neuron 86.3 (May 2015), pp. 800–812. issn: 0896-6273. doi: 10.1016/j.neuron.2015.03.024. url: http://dx.doi.org/10.1016/j.neuron.2015.03.024.

[53] Andrew J. Peters, Simon X. Chen, and Takaki Komiyama. “Emergence of reproducible spatiotemporal activity during motor learning.” In: Nature 510.7504 (May 2014), pp. 263–267. issn: 1476-4687. doi: 10.1038/nature13235. url: http://dx.doi.org/10.1038/nature13235.

[54] Woodrow L. Shew et al. “Neuronal Avalanches Imply Maximum Dynamic Range in Cortical Networks at Criticality.” In: The Journal of Neuroscience 29.49 (Dec. 2009), pp. 15595– 15600. issn: 1529-2401. doi: 10.1523/jneurosci.3864-09.2009. url: http://dx.doi.org/10.1523/JNEUROSCI.3864-09.2009.

[55] Joseph B. Kruskal. “Multidimensional scaling by optimizing goodness of fit to a nonmetric hypothesis.” In: Psychometrika 29.1 (Mar. 1964), pp. 1–27. issn: 1860-0980. doi: 10.1007/bf02289565. url: http://dx.doi.org/10.1007/BF02289565.

[56] Roger N. Shepard. “The analysis of proximities: Multidimensional scaling with an unknown distance function. I.” In: Psychometrika 27.2 (June 1962), pp. 125–140. issn: 1860-0980. doi: 10.1007/bf02289630. url: http://dx.doi.org/10.1007/BF02289630.

[57] Roger N. Shepard. “The analysis of proximities: Multidimensional scaling with an unknown distance function. II.” In: Psychometrika 27.3 (Sept. 1962), pp. 219–246. issn: 1860-0980. doi: 10.1007/bf02289621. url: http://dx.doi.org/10.1007/BF02289621.

[58] Jonathan D Cohen, Samuel M McClure, and Angela J Yu. “Should I stay or should I go? How the human brain manages the trade-off between exploitation and exploration.” In: Philosophical Transactions of the Royal Society B: Biological Sciences 362.1481 (Mar. 2007),pp. 933–942. issn: 1471-2970. doi: 10.1098/rstb.2007.2098. url: http://dx.doi.org/10.1098/rstb.2007.2098.

[59] L. P. Kaelbling, M. L. Littman, and A. W. Moore. “Reinforcement Learning: A Survey.” In: Journal of Artificial Intelligence Research 4 (May 1996), pp. 237–285. issn: 1076-9757. doi: 10.1613/jair.301. url: http://dx.doi.org/10.1613/jair.301.

[60] Neil J. Ritter, Nora M. Anderson, and Stephen D. Van Hooser. “Visual Stimulus Speed Does Not Influence the Rapid Emergence of Direction Selectivity in Ferret Visual Cortex.” In: The Journal of Neuroscience 37.6 (Jan. 2017), pp. 1557–1567. issn: 1529-2401. doi:10.1523/jneurosci.3365-16.2016. url: http://dx.doi.org/10.1523/JNEUROSCI.3365-16.2016.

[61] Torsten N. Wiesel and David H. Hubel. “SINGLE-CELL RESPONSES IN STRIATE CORTEX OF KITTENS DEPRIVED OF VISION IN ONE EYE.” In: Journal of Neurophysiology 26.6 (Nov. 1963), pp. 1003–1017. issn: 1522-1598. doi: 10.1152/jn.1963.26.6.1003. url: http://dx.doi.org/10.1152/jn.1963.26.6.1003.

[62] Ye Li et al. “Experience with moving visual stimuli drives the early development of cortical direction selectivity.” In: Nature 456.7224 (Oct. 2008), pp. 952–956. issn: 1476-4687. doi: 10.1038/nature07417. url: http://dx.doi.org/10.1038/nature07417.

[63] Deron Boyles and Jim Garrison. “The Mind Is Not the Brain: John Dewey, Neuroscience, and Avoiding the Mereological Fallacy.” In: Dewey Studies 1.1 (2017), pp. 111–130. url: https://opensiuc.lib.siu.edu/dewey-studies/vol1/iss1/23.

[64] Stuart S. Glennan. “Mechanisms and the nature of causation.” In: Erkenntnis 44.1 (Jan. 1996). issn: 1572-8420. doi: 10.1007/bf00172853. url: http://dx.doi.org/10.1007/BF00172853.

[65] Carl F. Craver. *Explaining the Brain: Mechanisms and the Mosaic Unity of Neuroscience*. Oxford: Oxford University Press, 2007. doi: 10.1093/acprof:oso/9780199299317.001. 0001.

[66] Marten Scheffer et al. “Early-warning signals for critical transitions.” In: Nature 461.7260 (Sept. 2009), pp. 53–59. issn: 1476-4687. doi: 10.1038/nature08227. url: http://dx.doi.org/10.1038/nature08227.

[67] Y. Bengio, P. Simard, and P. Frasconi. “Learning long-term dependencies with gradient descent is difficult.” In: IEEE Transactions on Neural Networks 5.2 (Mar. 1994), pp. 157–166. issn: 1941-0093. doi: 10.1109/72.279181. url: http://dx.doi.org/10.1109/72.279181.

[68] Razvan Pascanu, Tomas Mikolov, and Yoshua Bengio. “On the difficulty of training recurrent neural networks.” In: Proceedings of the 30th International Conference on Machine Learning. Ed. by Sanjoy Dasgupta and David McAllester. Vol. 28. Proceedings of Machine Learning Research 3. Atlanta, Georgia, USA: PMLR, 17–19 Jun 2013, pp. 1310–1318. url: https://proceedings.mlr.press/v28/pascanu13.html.

[69] Eric R. Kandel et al. Principles of Neural Science. 6th. New York: McGraw-Hill Education, 2021. isbn: 978-1259642234.

[70] Annika Hagemann et al. “Assessing criticality in pre-seizure single-neuron activity of human epileptic cortex.” In: PLOS Computational Biology 17.3 (Mar. 2021). Ed. by Peter Neal Taylor, e1008773. issn: 1553-7358. doi: 10.1371/journal.pcbi.1008773. url: https://dx.plos.org/10.1371/journal.pcbi.1008773.

[71] Ulric Neisser et al. “Intelligence: Knowns and unknowns.” In: American Psychologist 51.2 (1996), pp. 77–101. doi: 10.1037/0003-066X.51.2.77.

[72] Ian J. Deary, Alison Pattie, and John M. Starr. “The Stability of Intelligence From Age 11 to Age 90 Years: The Lothian Birth Cohort of 1921.” In: Psychological Science 24.12 (Oct. 2013), pp. 2361–2368. issn: 1467-9280. doi: 10.1177/0956797613486487. url: http://dx.doi.org/10.1177/0956797613486487.

73. Allen Institute for Brain Science. Allen Human Brain Atlas. Coronal Nissl reference atlas. 2010. url: https://human.brain-map.org (visited on 07/04/2026).

[74] Michael J. Hawrylycz et al. “An anatomically comprehensive atlas of the adult human brain transcriptome.” In: Nature 489.7416 (2012), pp. 391–399. doi: 10.1038/nature11405.

[75] Supratim Ray and John H. R. Maunsell. “Different Origins of Gamma Rhythm and High-Gamma Activity in Macaque Visual Cortex.” In: PLoS Biology 9.4 (Apr. 2011). Ed. by Leslie Ungerleider, e1000610. issn: 1545-7885. doi: 10.1371/journal.pbio.1000610. url:http://dx.doi.org/10.1371/journal.pbio.1000610.

[76] Jeremy R. Manning et al. “Broadband Shifts in Local Field Potential Power Spectra Are Correlated with Single-Neuron Spiking in Humans.” In: The Journal of Neuroscience 29.43 (Oct. 2009), pp. 13613–13620. issn: 1529-2401. doi: 10.1523/jneurosci.2041-09.2009. url: http://dx.doi.org/10.1523/JNEUROSCI.2041-09.2009.

[77] Longzhou Xu, Jianfeng Feng, and Lianchun Yu. “Avalanche criticality in individuals, fluid intelligence, and working memory.” In: Human Brain Mapping 43.8 (2022), pp. 2534–2553. issn: 10970193. doi: 10.1002/hbm.25802.

[78] Takahiro Ezaki et al. “Closer to critical resting-state neural dynamics in individuals with higher fluid intelligence.” In: Communications Biology 3.1 (2020). issn: 23993642. doi: 10.1038/s42003-020-0774-y. url: http://dx.doi.org/10.1038/s42003-020-0774-y.

[79] Gianina Cristian et al. “Critical Dynamics in the Association Cortex Predict Higher Intelligence in Typically Developing Children.” In: Journal of Neuroscience 46.9 (Mar. 2026), e1414252026. issn: 1529-2401. doi: 10.1523/JNEUROSCI.1414-25.2026. url: http://dx.doi.org/10.1523/JNEUROSCI.1414-25.2026.

[80] Renée A. Duckworth. “The role of behavior in evolution: a search for mechanism.” In: Evolutionary Ecology 23.4 (Mar. 2008), pp. 513–531. issn: 1573-8477. doi: 10.1007/s10682-008-9252-6. url: http://dx.doi.org/10.1007/s10682-008-9252-6.

[81] Craig V. Stewart and Dietmar Plenz. “Homeostasis of neuronal avalanches during postnatal cortex development in vitro.” In: Journal of Neuroscience Methods 169.2 (Apr. 2008), pp. 405–416. issn: 0165-0270. doi: 10.1016/j.jneumeth.2007.10.021. url: http\://dx.doi.org/10.1016/j.jneumeth.2007.10.021.

[82] James N. McGregor et al. “Failure in a population: Tauopathy disrupts homeostatic set-points in emergent dynamics despite stability in the constituent neurons.” In: Neuron 112.21 (Nov. 2024), 3567–3584.e5. issn: 0896-6273. doi: 10.1016/j.neuron.2024.08.006. url: http://dx.doi.org/10.1016/j.neuron.2024.08.006.

[83] Sarah L. Hissen et al. “The Stability and Repeatability of Spontaneous Sympathetic Baroreflex Sensitivity in Healthy Young Individuals.” In: Frontiers in Neuroscience 12 (2018). issn: 1662-453X. doi: 10.3389/fnins.2018.00403. url: http://dx.doi.org/10.3389/fnins.2018.00403.

[84] Christopher A. Sims. “Money, Income, and Causality.” In: The American Economic Review 62.4 (Sept. 1972), pp. 540–552.

[85] George Sugihara et al. “Detecting Causality in Complex Ecosystems.” In: Science 338.6106 (Oct. 2012), pp. 496–500. issn: 1095-9203. doi: 10.1126/science.1227079. url: http://dx.doi.org/10.1126/science.1227079.

[86] C. W. J. Granger. “Investigating Causal Relations by Econometric Models and Cross-spectral Methods.” In: Econometrica 37.3 (Aug. 1969), p. 424. issn: 0012-9682. doi: 10.2307/1912791. url: http://dx.doi.org/10.2307/1912791.

[87] David P White, et al. “Sleep deprivation and the control of ventilation.” In: Am. Rev. of Resp. Disease 128.6 (1983), pp. 984–986.

[88] William D.S. Killgore. “Effects of sleep deprivation on cognition.” In: Progress in Brain Research (2010), pp. 105–129.

[89] Christian Meisel et al. “Fading Signatures of Critical Brain Dynamics during Sustained Wakefulness in Humans.” In: The Journal of Neuroscience 33.44 (Oct. 2013), pp. 17363– 17372. issn: 1529-2401. doi: 10.1523/jneurosci.1516-13.2013. url: http://dx.doi.org/10.1523/JNEUROSCI.1516-13.2013.

[90] Chantel S. Prat et al. “Resting-state qEEG predicts rate of second language learning in adults.” In: Brain and Language 157–158 (2016), pp. 44–50. issn: 0093-934X. doi: 10.1016/j.bandl.2016.04.007. url: http://dx.doi.org/10.1016/j.bandl.2016.04.007.

[91] Maria Kliesch, Nathalie Giroud, and Martin Meyer. “EEG Resting-State and Event-Related Potentials as Markers of Learning Success in Older Adults Following Second Language Training: A Pilot Study.” In: Brain Plasticity 7.2 (Dec. 2021), pp. 143–162. issn: 2213-6312. doi: 10.3233/bpl-200117. url: http://dx.doi.org/10.3233/BPL-200117.

[92] Mianxin Liu et al. “Individual Cortical Entropy Profile: Test–Retest Reliability, Predictive Power for Cognitive Ability, and Neuroanatomical Foundation.” In: Cerebral Cortex Communications 1.1 (2020). issn: 2632-7376. doi: 10.1093/texcom/tgaa015. url: http://dx.doi.org/10.1093/texcom/tgaa015.

[93] Bence P Ölveczky, Aaron S Andalman, and Michale S Fee. “Vocal Experimentation in the Juvenile Songbird Requires a Basal Ganglia Circuit.” In: PLoS Biology 3.5 (Mar. 2005). Ed. by Wolfram Schultz, e153. issn: 1545-7885. doi: 10.1371/journal.pbio.0030153. url: http://dx.doi.org/10.1371/journal.pbio.0030153.

[94] Roxana Zeraati et al. “Neural timescales from a computational perspective.” In: Nature Neuroscience 29.7 (2026), pp. 1534–1547. issn: 1546-1726. doi: 10.1038/s41593-026-02343-8. url: http://dx.doi.org/10.1038/s41593-026-02343-8.

[95] Jaakko Matias Palva and Satu Palva. The Correlation of the Neuronal Long-Range Temporal Correlations, Avalanche Dynamics with the Behavioral Scaling Laws and Interindividual Variability. Mar. 2014. doi: 10.1002/9783527651009.ch5. url: http://dx.doi.org/10.1002/9783527651009.ch5.

[96] Jens Wilting et al. “Operating in a reverberating regime enables rapid tuning of network states to task requirements.” In: Frontiers in Systems Neuroscience 12 (Nov. 2018), p. 55. issn: 16625137. doi: 10.3389/fnsys.2018.00055. url: www.frontiersin.org.

[97] Nicholas M. Timme et al. “Criticality Maximizes Complexity in Neural Tissue.” In: Frontiers in Physiology 7 (2016). issn: 1664-042X. doi: 10.3389/fphys.2016.00425. url: http://dx.doi.org/10.3389/fphys.2016.00425.

[98] Anne-Sophie Darmaillacq, Ludovic Dickel, and Jennifer A Mather. Cephalopod cognition. Cambridge University Press, 2014.

[99] Marius Pachitariu et al. “A critical initialization for biological neural networks.” In: Nature (May 2026). issn: 1476-4687. doi: 10.1038/s41586-026-10528-1. url: http://dx.doi.org/10.1038/s41586-026-10528-1.

[100] Gina G. Turrigiano et al. “Activity-dependent scaling of quantal amplitude in neocortical neurons.” In: Nature 391.6670 (Feb. 1998), pp. 892–896. issn: 1476-4687. doi: 10.1038/36103. url: http://dx.doi.org/10.1038/36103.

[101] Ciaran Murphy-Royal, ShiNung Ching, and Thomas Papouin. “A conceptual framework for astrocyte function.” In: Nature Neuroscience 26.11 (Oct. 2023), pp. 1848–1856. issn: 1546-1726. doi: 10.1038/s41593-023-01448-8. url: http://dx.doi.org/10.1038/s41593-023-01448-8.

102. IEEE Standard for a Precision Clock Synchronization Protocol for Networked Measurement and Control Systems. IEEE Std 1588-2019 (Revision of IEEE Std 1588-2008). IEEE. New York, NY, USA, June 2020. doi: 10.1109/IEEESTD.2020.9120376. url: https://ieee.org.

[103] Jason E. Chung et al. “A Fully Automated Approach to Spike Sorting.” In: Neuron 95.6 (Sept. 2017), 1381–1394.e6. issn: 0896-6273. doi: 10.1016/j.neuron.2017.08.030. url: http://dx.doi.org/10.1016/j.neuron.2017.08.030.

[104] Alessio Paolo Buccino et al. “SpikeInterface, a unified framework for spike sorting.” In: eLife 9 (2020), e61834. doi: 10.7554/eLife.61834.

[105] Tianqi Chen and Carlos Guestrin. “XGBoost: A Scalable Tree Boosting System.” In: Proceedings of the 22nd ACM SIGKDD International Conference on Knowledge Discovery and Data Mining. KDD ’16. New York, NY, USA: ACM, 2016, pp. 785–794. doi: 10.1145/2939672.2939785.

[106] Peter Bartho et al. “Characterization of neocortical principal cells and interneurons by network interactions and extracellular features.” In: Journal of Neurophysiology 92.1 (2004), pp. 600–608. doi: 10.1152/jn.01170.2003.

[107] Alexander Mathis et al. “DeepLabCut: markerless pose estimation of user-defined body parts with deep learning.” In: Nature Neuroscience 21 (2018), pp. 1281–1289. doi: 10.1038/s41593-018-0209-y.

[108] Vladyslav V. Vyazovskiy et al. “Local sleep in awake rats.” In: Nature 472.7344 (Apr. 2011), pp. 443–447. issn: 1476-4687. doi: 10.1038/nature10009. url: http://dx.doi.org/10.1038/nature10009.

109. D. L. Ringach, R. M. Shapley, and M. J. Hawken. “Orientation selectivity in macaque V1: diversity and laminar dependence.” In: J Neurosci 22.13 (2002), pp. 5639–5651.

[110] M. Mazurek, M. Kager, and S. D. Van Hooser. “Robust quantification of orientation selectivity and direction selectivity.” In: Front Neural Circuits 8 (2014), p. 92. doi: 10.3389/fncir.2014.00092.

[111] Kenneth G. Wilson. “The renormalization group: Critical phenomena and the Kondo problem.” In: Rev. Mod. Phys. 47.4 (1975), p. 773. issn: 00346861. doi: 10.1103/RevModPhys.47.773.

[112] L. L. Thurstone. The Learning Curve Equation. Vol. 26. Psychological Monographs 3. Princeton, NJ: Psychological Review Company, 1919.

[113] Allen Newell and Paul S. Rosenbloom. “Mechanisms of Skill Acquisition and the Law of Practice.” In: Cognitive Skills and Their Acquisition. Ed. by John R. Anderson. Hillsdale, NJ: Lawrence Erlbaum Associates, 1981, pp. 1–55. isbn: 9780898590046.

[114] Paul M. Fitts and Michael I. Posner. Human Performance. Belmont, CA: Brooks/Cole, 1967.

[115] L. R. Rabiner. “A tutorial on hidden Markov models and selected applications in speech recognition.” In: Proceedings of the IEEE 77.2 (1989), pp. 257–286. doi: 10.1109/5.18626.

[116] hmmlearn developers. hmmlearn: Hidden Markov Models in Python. https://github.com/hmmlearn/hmmlearn. 2024.

[117] Carmelo Bellardita and Ole Kiehn. “Phenotypic Characterization of Speed-Associated Gait Changes in Mice Reveals Modular Organization of Locomotor Networks.” In: Current Biology 25.11 (2015), pp. 1426–1436. issn: 0960-9822. doi: 10.1016/j.cub.2015.04.005. url: http://dx.doi.org/10.1016/j.cub.2015.04.005.

[118] Ingwer Borg and Patrick J. F. Groenen. Modern Multidimensional Scaling: Theory and Applications. 2nd. New York, NY: Springer, 2005. doi: 10.1007/0-387-28981-X.

[119] S. B. Needleman and C. D. Wunsch. “A general method applicable to the search for similarities in the amino acid sequence of two proteins.” In: Journal of Molecular Biology 48.3 (1970), pp. 443–453. doi: 10.1016/0022-2836(70)90057-4.

[120] Daniel Abásolo, et al. “Lempel-Ziv complexity of cortical activity during sleep and waking in rats.” In: Journal of Neurophysiology 113.7 (2015), pp. 2742–2752. doi: 10.1152/jn.00575.2014.

[121] Leandro J. Fosque et al. “Evidence for Quasicritical Brain Dynamics.” In: Physical Review Letters 126.9 (Mar. 2021). issn: 1079-7114. doi: 10.1103/physrevlett.126.098101. url: http://dx.doi.org/10.1103/PhysRevLett.126.098101.

